# A molecular switch orchestrates the nuclear export of human messenger RNA

**DOI:** 10.1101/2024.03.24.586400

**Authors:** Ulrich Hohmann, Max Graf, Ulla Schellhaas, Belén Pacheco-Fiallos, Laura Fin, Daria Riabov-Bassat, Thomas Pühringer, Michael-Florian Szalay, László Tirián, Dominik Handler, Julius Brennecke, Clemens Plaschka

## Abstract

The nuclear export of messenger RNA (mRNA) is a key step in eukaryotic gene expression (Köhler and Hurt, 2007). Despite recent insights into the packaging of newly transcribed mRNAs into ribonucleoprotein complexes (mRNPs) (Pacheco-Fiallos et al., 2023; Bonneau et al., 2023), the subsequent events that govern mRNA export are poorly understood. Here, we elucidate the molecular basis of human mRNA export licensing, which involves the remodeling of mRNP-bound transcription-export complexes (TREX), the formation of export-competent mRNPs, the docking of mRNPs at the nuclear pore complex (NPC), and the release of mRNPs at the NPC to initiate export. Our biochemical and structural data uncover the ATPase DDX39/UAP56 as a central molecular switch that directs mRNPs through the TREX and the NPC-anchored TREX-2 complexes using its ATPase and mRNA-binding cycle. Collectively, these findings establish a mechanistic framework for a general and conserved mRNA export pathway.

## INTRODUCTION

Eukaryotic gene expression requires the nuclear export of newly synthesized messenger RNAs (mRNA) through the nuclear pore complex (NPC) for translation in the cytoplasm. To prevent the translation of aberrant RNAs, mRNA export is selective for mature mRNA ribonucleoprotein complexes (mRNPs).

Mature mRNPs are marked by specific proteins, which they acquire during the capping, splicing, cleavage, and polyadenylation of their precursor mRNAs (Köhler and Hurt, 2007; Singh et al., 2015; Heath et al., 2016; Khong and Parker, 2020; Vorländer et al., 2022). By recognizing these maturation marks, the transcription-export complex (TREX) assembles on the surface of packaged mRNPs and selects maturing mRNAs for export (Cheng et al., 2006; Köhler and Hurt, 2007; Kelly and Corbett, 2009; Heath et al., 2016; Pacheco-Fiallos et al., 2023). TREX also aids in mRNA packaging and thereby ensures genome integrity by preventing the formation of harmful RNA–DNA hybrids, called R-loops (Huertas and Aguilera, 2003). However, packaged TREX–mRNP complexes cannot be directly exported (Dias et al., 2010; Chi et al., 2013). Instead, they need to undergo a two-step ‘licensing process’. First, TREX must be disassembled to generate export-competent mRNPs (Köhler and Hurt, 2007). Second, these remodeled mRNPs must engage the NPC, where the global mRNA export factor, NXF1–NXT1, facilitates mRNP transport across the NPC’s selective permeability barrier (Gatfield et al., 2001; Sträßer et al., 2002; Zenklusen et al., 2002; Viphakone et al., 2012; Lee et al., 2020). While the factors required for mRNA export were discovered decades ago (Luo et al., 2001; Sträßer and Hurt, 2001; Sträßer et al., 2002; Fischer et al., 2002), the mechanistic basis of the different steps underlying mRNA export licensing and how mRNPs navigate through these steps remains unclear.

Using a combination of biochemistry, in silico protein-protein interaction screening, cryo-electron microscopy, and cellular assays, we here elucidate a general mechanism for mRNA export that assigns molecular functions to key mRNA export proteins and their complexes.

## RESULTS

### UAP56 bridges the THO complex to mRNPs

Nuclear mRNPs form compact globules (Skoglund et al., 1986; Pacheco-Fiallos et al., 2023; Bonneau et al., 2023) decorated with TREX complexes on their surface (Pacheco-Fiallos et al., 2023) (Fig. 1a). To investigate how TREX–mRNP complexes are subsequently remodeled for nuclear export, we re-examined how TREX interacts with mRNPs after their recognition and packaging. In humans, TREX comprises a tetramer of the 6-subunit THO complex, each containing THOC1, −2, −3, −5, −6, −7, four DDX39/UAP56 (UAP56 hereafter; yeast Sub2) subunits, and various mRNA export adaptors such as ALYREF (yeast Yra1) (Sträßer et al., 2002; Heath et al., 2016). ALYREF interacts directly with mRNP-bound maturation marks, such as the exon junction complex (EJC) or the cap binding complex. ALYREF also binds to UAP56 through N- and C-terminal UAP56-binding motifs (N- and C-UBM) (Le Hir et al., 2000; Sträßer et al., 2002; Heath et al., 2016), though only the C-UBM had been observed in structures (Ren et al., 2017; Pacheco-Fiallos et al., 2023) (Fig. 1d). UAP56 is a DExD-box ATPase whose two RecA lobes, RecA1 and RecA2, can stably clamp onto RNA together with ATP (Ren et al., 2017) (Fig. 1d,e). In the cryo-EM structures of native TREX–mRNP complexes (Pacheco-Fiallos et al., 2023), the four UAP56 molecules are however not yet clamped onto RNA (Fig. 1b-e). Instead, UAP56 is ‘primed’ for mRNA-clamping. The RecA2 lobe of each UAP56 interacts with one of the four THOC2 subunits (Ren et al., 2017; Pühringer et al., 2020; Schuller et al., 2020; Xie et al., 2021) (Fig. 1b,c), whereas each RecA1 lobe bridges to the mRNP through an interaction with the C-terminal UBM (C-UBM) in ALYREF (Hautbergue et al., 2009; Gromadzka et al., 2016; Pacheco-Fiallos et al., 2023) (Fig. 1b-d).

**Figure 1.**
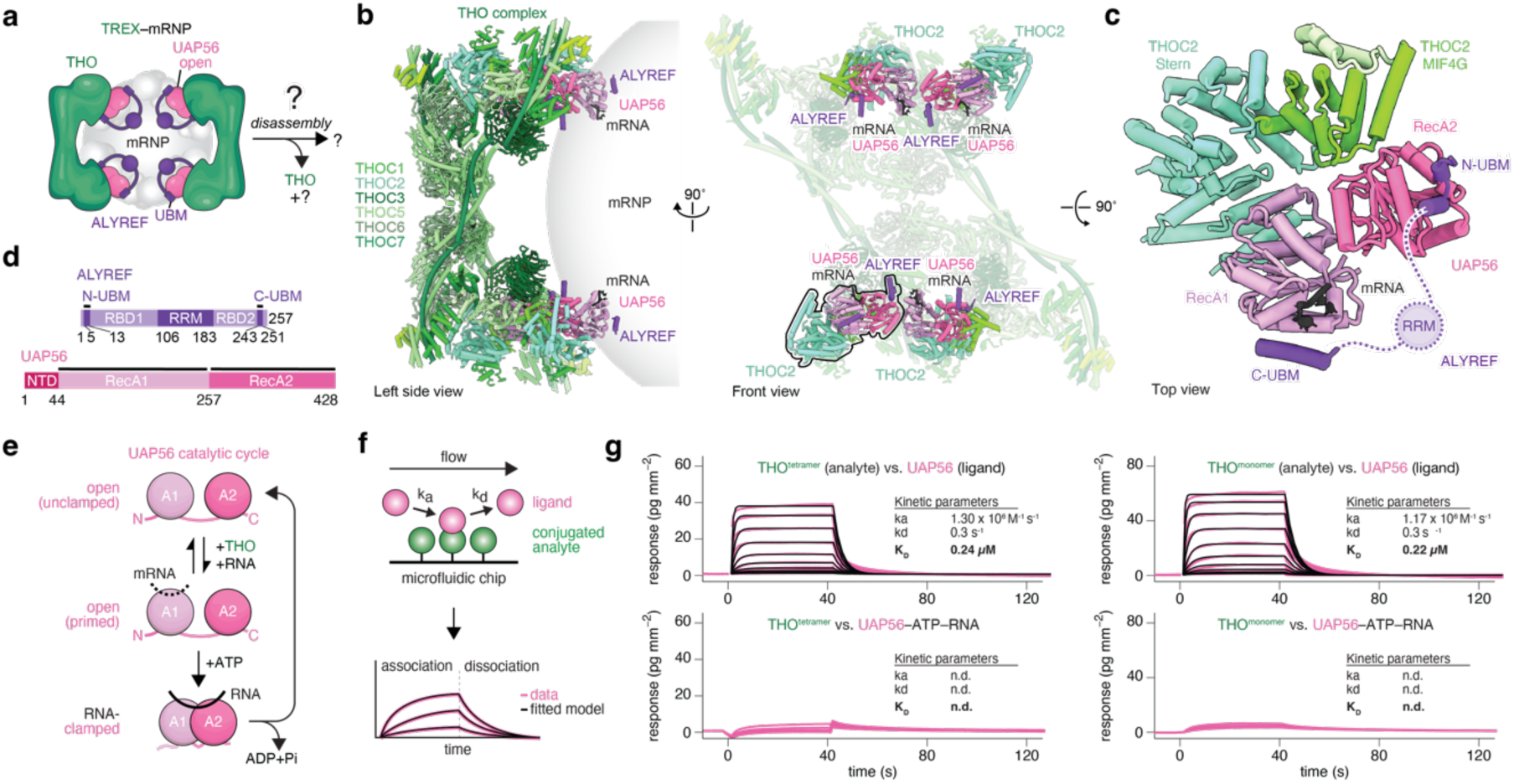
UAP56 controls TREX–mRNP assembly and disassembly. **a,** Cartoon schematic of a TREX–mRNP complex, which prior to mRNP export must disassemble TREX and remodel the mRNP via an unknown mechanism. **b-c,** Newly revised cryo-EM structure of a human TREX–mRNA complex shown from the left side and the front (left and middle panels) with **c,** a closeup of the TREX–mRNA interface shown from the top (right panel). UAP56 binds the THO subunit THOC2 MIF4G and Stern domains, and is in an open conformation, primed to clamp onto the associated segment of putative mRNA (black). UAP56 further binds ALYREF through an N- and C-UBM at its respective RecA2 and RecA2 lobes (also see Supplementary Fig. 2g). The UAP56 N-terminal domain is disordered. THO, shades of green; UAP56, shades of pink; ALYREF, purple; mRNP, grey sphere. UBM, UAP56-binding motif. **d,** Domain organization of ALYREF and UAP56. Regions included in the atomic model are indicated with a black line. RBD, RNA binding domain; RRM, RNA Recognition Motif; NTD, N-terminal domain. **e,** Cartoon schematic of the UAP56 catalytic cycle. In the open state, the UAP56 RecA1 and RecA2 lobes are not coordinated (top). Binding to the THO subunit THOC2 induces a primed state, where RNA may associate with the RecA1 lobe, but UAP56 stays open (middle). In the closed state, UAP56 binds ATP and clamps onto RNA by coordinating the RecA1 and RecA2 lobes (bottom). ATP hydrolysis and RNA release revert UAP56 from the RNA-clamped to an unclamped, open state. **f,** Experiment scheme for Grating-coupled interferometry (GCI). The analyte is immobilized on a microfluidic chip and a putative ligand is flown in at increasing concentrations (association), and subsequently washed out with buffer (dissociation). Binding is recorded as a change in the refractive index, yielding ‘sensograms’, which are colored by the respective ligand and fitted with a 1-to-1 binding kinetic model (overlayed black line). See Methods for details. **g,** Sensograms for tetrameric or monomeric THO (immobilized) probed with UAP56 or UAP56–ATP–RNA. Sensograms (pink line), the fitted model (black), and a binding kinetics summary table are shown. Isolated, open UAP56 binds the tetrameric or monomeric THO complex with comparable affinities (top row), whereas RNA-clamped, closed UAP56–ATP– RNA has no measurable THO complex affinity (bottom row).

ALYREF contains a second UBM at its N-terminus (N-UBM) (Golovanov et al., 2006; Hautbergue et al., 2009). While it has been proposed that the N-UBM mimics the C-UBM in binding to the RecA1 lobe of UAP56 (Hautbergue et al., 2009; Gromadzka et al., 2016; Pühringer et al., 2020; Schuller et al., 2020), the amino acid sequences of the two UBMs differ despite each being highly conserved. When we re-examined published TREX–mRNP maps (Pacheco-Fiallos et al., 2023), we noticed a low-resolution density that was consistent with an AlphFold2 prediction of the ALYREF N-UBM bound to the UAP56 RecA2 lobe (Jumper et al., 2021; Evans et al., 2022; Mirdita et al., 2022) (Fig. 1b, c, Supplementary Fig. 1a-e). To test this prediction, we reconstituted a TREX–mRNP complex *in vitro* comprising the recombinant human THO complex, UAP56, the EJC, a 15 nucleotide single-stranded poly-Uridine RNA (15U RNA), and an ALYREF construct containing only the N-UBM, N-terminal arginine-rich linker and RRM, but lacking the C-terminal arginine-rich linker and C-UBM (ALYREF residues 1-183) (Pacheco-Fiallos et al., 2023) (Supplementary Fig. 2a). Cryo-EM analysis of this complex revealed the THO–UAP56 protomer at 4.1 Å resolution, allowing for the unambiguous assignment of the ALYREF N-UBM on the UAP56 RecA2 lobe (Supplementary Fig. 2b-g; Supplementary Table 1). Consistent with two distinct UBM binding sites, mutating either of these binding sites in UAP56 abolished the N-UBM peptide or C-UBM peptide interaction, respectively (Supplementary Fig. 1f).

Thus, ALYREF binds to two distinct sites on UAP56, forming unique composite surfaces that could be leveraged for mRNA export (see below). Furthermore, UAP56 stands out as the sole protein identified to bridge the THO complex and ALYREF-bound mRNPs (Fig. 1b,c). In this assembly, UAP56 assumes a ‘primed’ conformation, poised to clamp onto mRNA (Pacheco-Fiallos et al., 2023).

### RNA-clamping releases UAP56 from the THO complex

To facilitate the nuclear export of mRNPs, TREX disassembles by an unknown mechanism (Köhler and Hurt, 2007; Dias et al., 2010; Chi et al., 2013). Given the central position of UAP56 in TREX as a bridge between THO and the mRNP, we explored whether the ATP-dependent mRNA-clamping of UAP56 and the ensuing conformational change (Ren et al., 2017) might serve as a molecular mechanism to render UAP56 incompatible with THO-binding. To test this, we used Grating Coupled Interferometry (GCI) (Kozma et al., 2009) and measured the affinity of a surface-immobilized THO complex for either UAP56 alone or for ATP-bound, RNA-clamped UAP56 (Fig. 1f,g). This showed that the THO complex binds isolated UAP56 with a K_D_ of approximately 0.25 µM. In contrast, THO exhibited no measurable binding affinity for the UAP56–ATP–RNA complex (Fig. 1g). Unexpectedly however, when we pre-formed THO– UAP56 complexes using recombinant proteins, we observed only partial complex disassembly upon the addition of ATP and RNA (Supplementary Fig. 3b). Given that the disassembly of TREX necessitates the coordinated release of all four THO-bound UAP56 molecules and no cooperative binding was observed between UAP56 and tetrameric, dimeric, or monomeric THO complexes (Fig. 1g, Supplementary Fig. 3a), we hypothesized that a dedicated factor, in addition to RNA-clamping by UAP56, may be required for the efficient disassembly of multivalent TREX–mRNP complexes (Fig. 2a).

**Figure 2.**
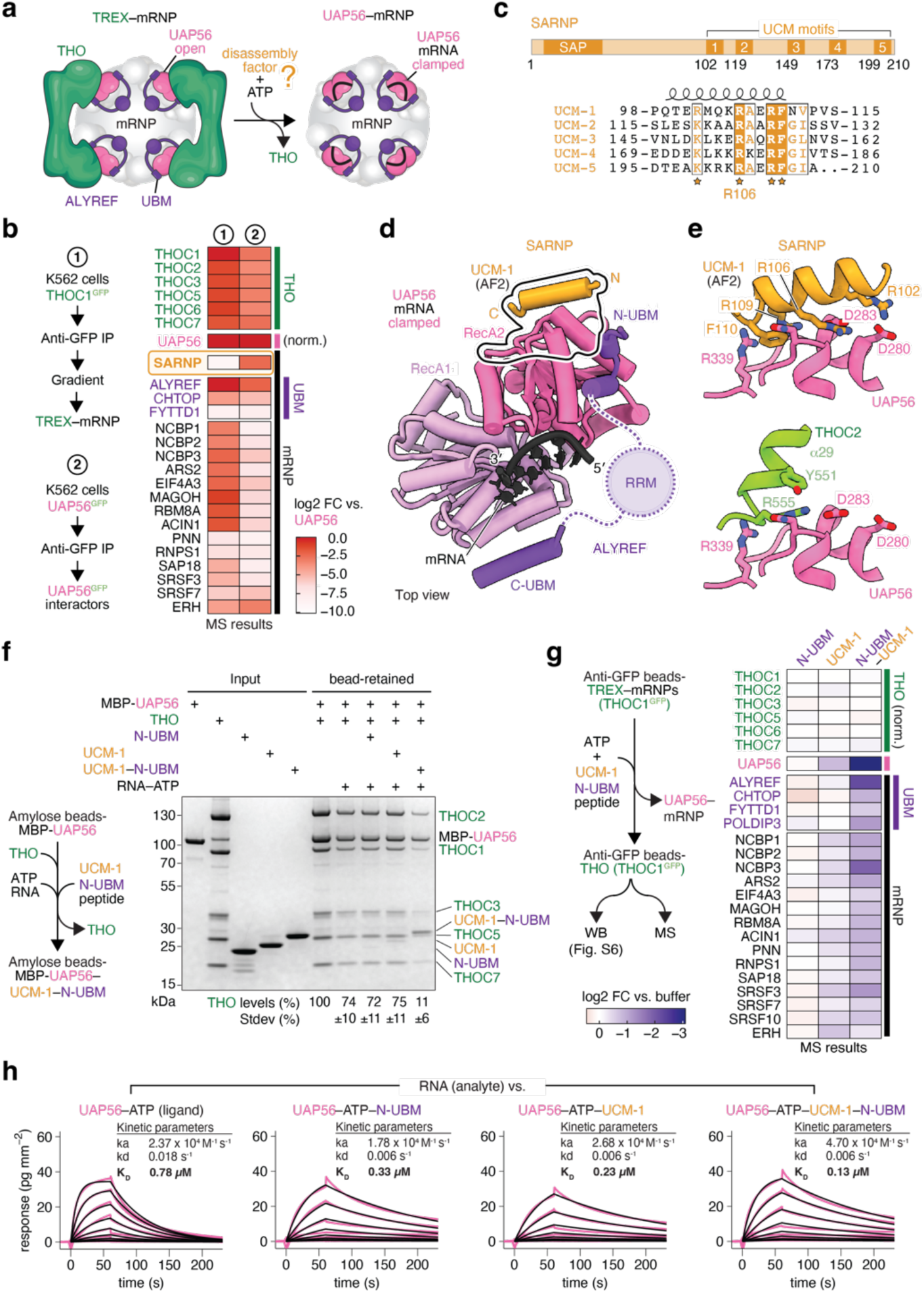
SARNP assists TREX–mRNP disassembly and UAP56–mRNA stabilization. **a,** Cartoon schematic for TREX–mRNP disassembly, which may require an additional disassembly factor for efficient remodeling. **b,** Heatmap comparing the abundance of mRNP-associated proteins in purified TREX–mRNPs (Pacheco-Fiallos et al., 2023) versus the nuclear UAP56 protein interactome, as determined by mass spectrometry. The heatmap is colored according to the log2 fold-change after normalizing to UAP56 levels. The UAP56 protein interactome was determined from a GFP-UAP56 immunoprecipitation in human K562 cell nuclear extract (see Methods). SARNP was absent in TREX–mRNPs, but abundant in the UAP56 protein interactome. **c,** Domain organization (top) and multiple sequence alignment (bottom) of human SARNP (Uniprot ID P82979) and its five UAP56-clamping motifs (UCMs). Residues invariant or conserved among the five UCMs (UCM-1 to −5) are highlighted in orange. **d,** Model of a clamped UAP56–ATP–RNA complex bound to SARNP and ALYREF. The model was obtained by superposing structures of clamped UAP56–ATP–RNA (PDB-ID 8ENK) (Xie et al., 2023), open UAP56–ALYREF N- and C-UBMs (Fig. 1), and a UAP56– SARNP UCM-1 AlphaFold2 Multimer prediction (Supplementary Figs. 1b-e, 4a-d). UAP56, pink; RNA, black; SARNP, orange; ALYREF, purple. UBM, UAP56-binding motif. **e,** Details of the UAP56–SARNP UCM-1 model with key interface residues shown as sticks (top). Shown below is the same view of UAP56 in the THO–UAP56 complex (THOC2, green) (Pühringer et al., 2020), revealing a steric clash between SARNP and THOC2. Colors as in panel **d**. **f,** THO–UAP56 disassembly assay with recombinant proteins. Experiment schematic (left) and SDS-PAGE analysis of the results are shown (right, Coomassie stain). The amount of bead-retained THO complex is quantified underneath from three independent experiments. **g,** TREX–mRNP disassembly assay with native complexes. Experiment schematic (left) and mass spectrometry results (right, heatmap) of bead-retained mRNP-associated proteins after adding the ALYREF N-UBM, SARNP UCM-1, or an UCM-1–N-UBM fusion. The heatmap is colored according to the log2 fold-change compared to the buffer control, after normalizing to mean THO complex subunit levels. See Methods for details and a western blot analysis in Supplementary Fig. 5b. **h,** Sensograms for a 15 nucleotide poly-Uridine RNA (immobilized) probed with UAP56, UAP56–UCM-1, UAP56–N-UBM or an UAP56–UCM-1–N-UBM fusion, with ATP in all buffers. Sensograms (pink line), the fitted model (black), and a binding kinetics summary table are shown, revealing that N-UBM and UCM-1 increase the affinity of UAP56–ATP to RNA approximately 6-fold.

### SARNP is a multivalent TREX disassembly factor

To identify a factor contributing to TREX disassembly, we compared the protein composition of TREX-bound mRNPs (Pacheco-Fiallos et al., 2023) with the nuclear UAP56 protein interactome in human cells (Fig. 2b, Supplementary Table 2). This analysis revealed a single protein, SARNP (CIP29, yeast Tho1), that was absent from purified TREX–mRNPs but highly enriched in the UAP56 interactome. Notably, SARNP has been implicated in mRNA export in yeast (Piruat and Aguilera, 1998; Jimeno et al., 2006), is conserved from yeast to humans, is essential for viability in human cells (Dempster et al., 2019), and is required for human mRNA export (Germain et al., 2010; Kang et al., 2020; Xie et al., 2023). Additionally, SARNP has been shown to bind human mRNAs *in vivo* (Baltz et al., 2012; Castello et al., 2012) and RNA-clamped UAP56 *in vitro* (Dufu et al., 2010; Xie et al., 2023) in the absence of the THO complex (Sørensen et al., 2017; Bonneau et al., 2023), consistent with the anticipated activities of a TREX disassembly factor.

Using AlphaFold2 (Jumper et al., 2021; Evans et al., 2022; Mirdita et al., 2022), we predicted a direct interaction between human UAP56 and human SARNP (Fig. 2c-e; Supplementary Fig. 4a-d). This prediction was indistinguishable from a recent crystal structure of a chimeric human UAP56–yeast SARNP complex exhibiting an RMSD of 0.67 Å across 370 atom pairs (Xie et al., 2023). In these models, an α-helical motif within SARNP binds to the UAP56 RecA2 lobe (Fig. 2d,e). Mutations of the highly conserved residue R106D in the SARNP motif (residues 81-115) or of D283R in the UAP56 RecA2 lobe disrupted the UAP56–SARNP motif interaction *in vitro* (Fig. 2e, Supplementary Fig. 4e,f). Due to its distinctive biochemical activities described below, we refer to SARNP’s UAP56 interacting peptide as the ‘<u>U</u>AP56 <u>c</u>lamping <u>m</u>otif’ (UCM). The UCM is found five times in human SARNP with the consensus sequence R/KxxxRAxRFG (Fig. 2c, Supplementary Fig. 4g).

The binding site for the SARNP UCM on the UAP56 RecA2 lobe is located directly adjacent to the newly identified binding site for the ALYREF N-UBM (Figs. 1b, 2d). To explore whether UCM and N-UBM motifs bind cooperatively to UAP56, we first determined the affinities between UAP56 and isolated UCM (K_D_=12 µM) or N-UBM peptides (K_D_=24 µM) (Supplementary Figs. 1f, 4h). To address the challenges posed by such low-affinity complexes, we generated different ‘single-chain fusions’ with UAP56, a common strategy to stabilize low affinity interactions (Bird et al., 1988; Ming et al., 2019). Specifically, we fused UAP56 with either the SARNP UCM-1, the ALYREF N-UBM, or both. *In vitro* competition experiments with N-UBM or UCM peptides confirmed that the fused counterparts bound their cognate UAP56- binding sites (Supplementary Fig. 5a). When we then tested for cooperative binding, we indeed observed enhanced binding of the ALYREF N-UBM to the UAP56–UCM fusion (Supplementary Fig. 5a).

Notably, binding of the SARNP UCM to UAP56 would sterically clash with the interaction between UAP56 and the THOC2 MIF4G domain in TREX–mRNPs (Fig. 2e, Supplementary Fig. 4i). Consequently, UCM-binding to UAP56 might prevent the re-binding of UAP56 to THOC2, after UAP56 has clamped onto mRNA. Therefore, SARNP could promote TREX disassembly and directionality of these steps, assigning a definite molecular activity to this protein. To test this model, we reconstituted the THO–UAP56 complex on beads and examined its integrity upon addition of purified proteins in the presence of ATP and RNA. The addition of ATP and RNA alone or together with either an ALYREF N-UBM peptide or a SARNP UCM-1 peptide resulted in the comparable, but only partial, dissociation of THO from UAP56 (Fig. 2f, Supplementary Fig. 3b). Instead, the THO–UAP56 complex disassembled almost completely when we added a peptide comprising the SARNP UCM fused to the ALYREF N-UBM (Fig. 2f). These data suggest that cooperative binding of the SARNP UCM with an N-UBM, RNA, and ATP to UAP56 promotes the efficient dissociation of the THO–UAP56 interaction and thus TREX disassembly.

To challenge this model in a more native setting, we assessed whether SARNP and mRNA-clamping by UAP56 could disassemble TREX on endogenous mRNPs. We immobilized TREX–mRNPs purified from nuclear extract of human K562 cells on beads via the GFP-tagged THOC1 subunit (Pühringer et al., 2020) and added ATP, or ATP together with either the ALYREF N-UBM, the SARNP UCM, or a UCM–N-UBM fusion peptide (Fig. 2g). Upon disassembly, UAP56–mRNPs would be released from THO and thus depleted from the beads while bead-bound THO complexes would remain. We measured mRNP depletion using mass spectrometry and western blotting (Fig. 2g, Supplementary Fig. 5b, Supplementary Table 3). Addition of the N-UBM alone did not result in mRNP release (Fig. 2g, Supplementary Fig. 5b). In contrast, addition of the SARNP UCM resulted in detectable mRNP release, possibly aided by endogenous ALYREF that co-purified with mRNPs (Pacheco-Fiallos et al., 2023). This effect was further enhanced by the addition of the N-UBM–UCM fusion, which would bind to mRNP-bound UAP56 molecules cooperatively (Fig. 2g, Supplementary Fig. 5b). Collectively, these data support a role for UAP56 as the central bridge between the mRNP and the THO complex and demonstrate that the ATP-dependent mRNA-clamping of UAP56, assisted by ALYREF and SARNP, is sufficient to disassemble TREX. Human SARNP contains five UCMs connected by low-complexity linkers (Fig. 2c). We speculate that this UCM multivalency may have evolved to disassemble the multivalent tetrameric THO–UAP56 complexes in native mRNPs.

### SARNP stabilizes UAP56–RNA complexes

To prevent re-association of the mRNP with THO, UAP56 must stably clamp onto mRNA. We hypothesized that SARNP and ALYREF may stabilize RNA-clamped UAP56. Indeed, we observed that SARNP, as well as the fusion of ALYREF N-UBM and SARNP UCM to UAP56, enhanced the UAP56–RNA interaction *in vitro* (Supplementary Fig. 5c). To quantify this effect, we measured the RNA affinity of UAP56, UAP56–N-UBM, UAP56–UCM and UAP56–UCM– N-UBM. The N-UBM fusion increased UAP56’s affinity to RNA by approximately 2-fold (K_D_=0.78 µM vs. K_D_=0.33 µM), while the SARNP UCM fusion increased it by approximately 3.5-fold (K_D_=0.23 µM) (Fig. 2h). The UAP56–UCM–N-UBM fusion displayed a ∼6-fold higher RNA affinity (K_D_=0.13 µM) (Fig. 2h). Thus, SARNP and UAP56 mRNA-clamping do not only promote THO release but also stabilize UAP56–mRNP complexes. RNA-clamped UAP56 could therefore determine the downstream fate of the mRNP, such as their docking at the NPC (Fig. 3a).

**Figure 3.**
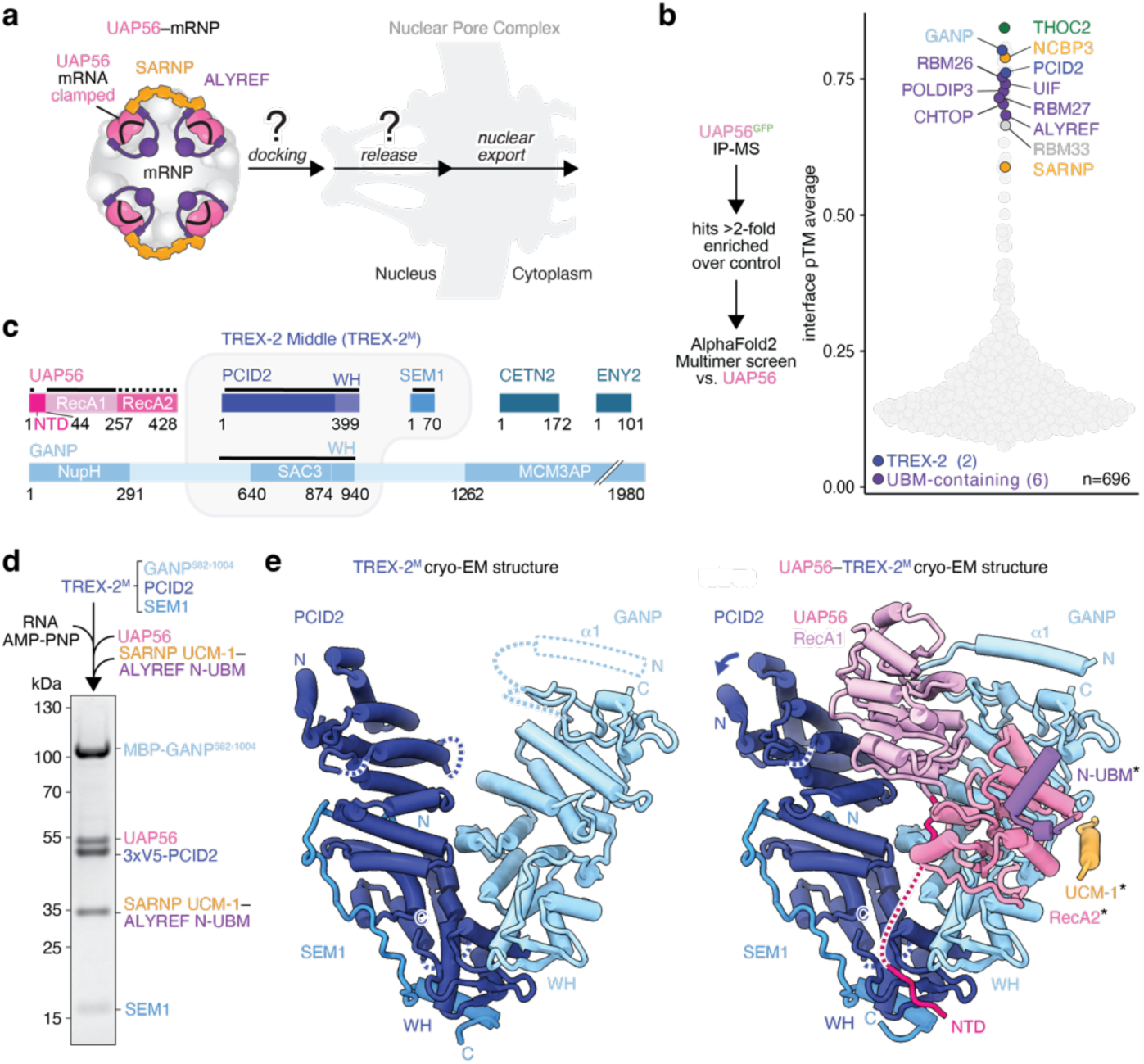
UAP56 binds the NPC-anchored TREX-2 complex. **a,** Cartoon schematic of a UAP56–mRNP complex, which docks at the NPC through an unknown mechanism prior to export. **b,** AlphaFold2 Multimer (Jumper et al., 2021; Evans et al., 2022; Mirdita et al., 2022) *in silico* protein interaction screen to identify novel UAP56 interactors. An experiment schematic (left) and the screen results (right) are shown. All UAP56–candidate predictions are shown (circles) and ranked by the average interface prediction TM score (interface pTM). Known and several novel predicted interactors are highlighted in colors. **c,** Domain organization of UAP56 (pink) and TREX-2 complex subunits (blue). Regions included in the atomic model are indicated with a black line, whereas rigid-body fits are indicated with a dotted line. WH, Winged Helix; NupH, Nucleoporin Homology. **d,** Reconstitution of a recombinant TREX-2^M^ complex with an UAP56–UCM-1–N-UBM assembly (see Supplementary Fig. 7a for details). SDS-PAGE analysis (Coomassie stain) of a representative *in vitro* protein pulldown is shown. **e,** Cryo-EM structures of human TREX-2^M^ (left) and UAP56–TREX-2^M^ (right) shown from the front. The UAP56 RecA2 lobe, UCM-1, and N-UBM are putatively fitted based on a low-resolution density (see Supplementary Fig. 8 for details). SEM1, blue; PCID2, dark blue; GANP, light blue; UAP56 shades of pink; ALYREF N-UBM, purple; SARNP UCM-1, orange.

### UAP56–RNA binds the NPC-anchored TREX-2 complex

Single molecule tracking experiments of mRNAs in yeast and human cells revealed that mRNPs transiently dock at NPCs prior to their nuclear export (Grünwald and Singer, 2010; Mor et al., 2010; Ma et al., 2013; Smith et al., 2015). The molecular mechanism underlying these events is not known (Fig. 3a). To identify proteins that might engage with UAP56–mRNPs after TREX disassembly, we first generated a comprehensive list of putative UAP56 interactors based on their greater than 2-fold enrichment in UAP56 immuno-precipitates from K562 cell nuclear extract. We then performed a pairwise AlphaFold2 Multimer (Jumper et al., 2021; Evans et al., 2022; Mirdita et al., 2022) protein interaction screen between each of these candidates and UAP56, and ranked the results by their interface pTM (ipTM) scores (Fig. 3b, Supplementary Table 4). Top scoring candidates included known UAP56 interactors, such as THOC2, SARNP, and the N- or C-UBM containing export adaptors ALYREF, CHTOP, LUZP4, UIF, and PHAX (Gromadzka et al., 2016). Several other proteins with known roles in nuclear mRNA metabolism, including RBM26, RBM27 and NCBP3, emerged as putative UAP56 interactors (Gebhardt et al., 2015; Dou et al., 2020; Silla et al., 2020) (Supplementary Fig. 6).

Among the top-ranking predicted UAP56 interactors were also GANP and PCID2, which are two of the five subunits of the NPC-anchored TREX-2 complex (Fischer et al., 2002, 2004; Rodríguez-Navarro et al., 2004; Wilmes et al., 2008; Faza et al., 2009; Jani et al., 2012) (Fig. 3b,c). TREX-2 is required for mRNA export, but its molecular functions are unclear (Fischer et al., 2002; Rondón et al., 2010; Wickramasinghe et al., 2010). The human TREX-2 complex consists of GANP, PCID2, SEM1, ENY2, and CETN2 or CETN3 (yeast Sac3, Thp1, Dss1, Cdc31 and Sus1, respectively) (Fischer et al., 2002, 2004; Rodríguez-Navarro et al., 2004; Wilmes et al., 2008; Faza et al., 2009; Jani et al., 2012). The GANP subunit serves as a scaffold for the four other subunits and anchors TREX-2 to the nuclear basket of the NPC (Fischer et al., 2002; Kurshakova et al., 2007; Jani et al., 2009; Lu et al., 2010). While TREX-2 has been reported to bind the mRNA export factor NXF1–NXT1 (Fischer et al., 2002; Dimitrova et al., 2015), no direct link to UAP56 has been described. The predicted interaction between UAP56 and TREX-2 therefore suggested an intriguing model in which TREX and TREX-2 act in a linear pathway where UAP56–mRNPs, after their release from THO, dock at the NPC via TREX-2 to facilitate the loading of NXF1–NXT1.

To investigate whether UAP56 binds to GANP and PCID2, we purified a minimal recombinant TREX-2 complex (TREX-2^M^) comprising the GANP Sac3 domain (residues 582-1004), PCID2, and SEM1 (Fig. 3c). In *in vitro* pulldown experiments, TREX-2^M^ exhibited a stochiometric interaction with UAP56 or a UAP56–UCM–N-UBM complex bound to RNA and AMP-PNP, confirming their direct interaction (Fig. 3d, Supplementary Fig. 7a).

To reveal the molecular interfaces underlying the interaction of UAP56 with TREX-2, we obtained cryo-EM structures of the TREX-2^M^ complex both in isolation (3.5 Å resolution) and bound to a UAP56–UCM–N-UBM complex (3.5 Å resolution) (Fig. 3e, Supplementary Figs. 7b-f, 8, Supplementary Table 1). The cryo-EM structure of the human TREX-2^M^ complex in isolation is consistent with reported structures of the yeast TREX-2^M^ complex (Ellisdon et al., 2012; Schneider et al., 2015; Aibara et al., 2016; Gordon et al., 2017) (Supplementary Fig. 9a,b), and exhibits a V-shaped architecture formed by the GANP Sac3 domain and PCID2 (Fig. 3e). SEM1 is largely unstructured and binds to PCID2 (Fig. 3e). In the novel UAP56–TREX-2^M^ structure, the N-terminal half of PCID2 rotates slightly outwards, hinging at residues 202-205, to accommodate UAP56, and an additional loop and helix in the GANP N-terminus become ordered compared to the apo state (Fig. 3e). Notably, although we performed these experiments with RNA-bound UAP56, the cryo-EM structure indicates that UAP56 is in an open conformation, not bound to RNA. While the UAP56 RecA1 lobe is resolved and binds between the ‘V’, formed by GANP and PCID2, the RecA2 lobe is mobile and poorly resolved (Fig. 3e, Supplementary Fig. 8d). These findings suggest that UAP56 facilitates the docking of its bound mRNPs at the NPC through interactions with TREX-2, and that TREX-2 may subsequently affect UAP56–mRNP complexes (see below) (Fig. 4a,b).

**Figure 4.**
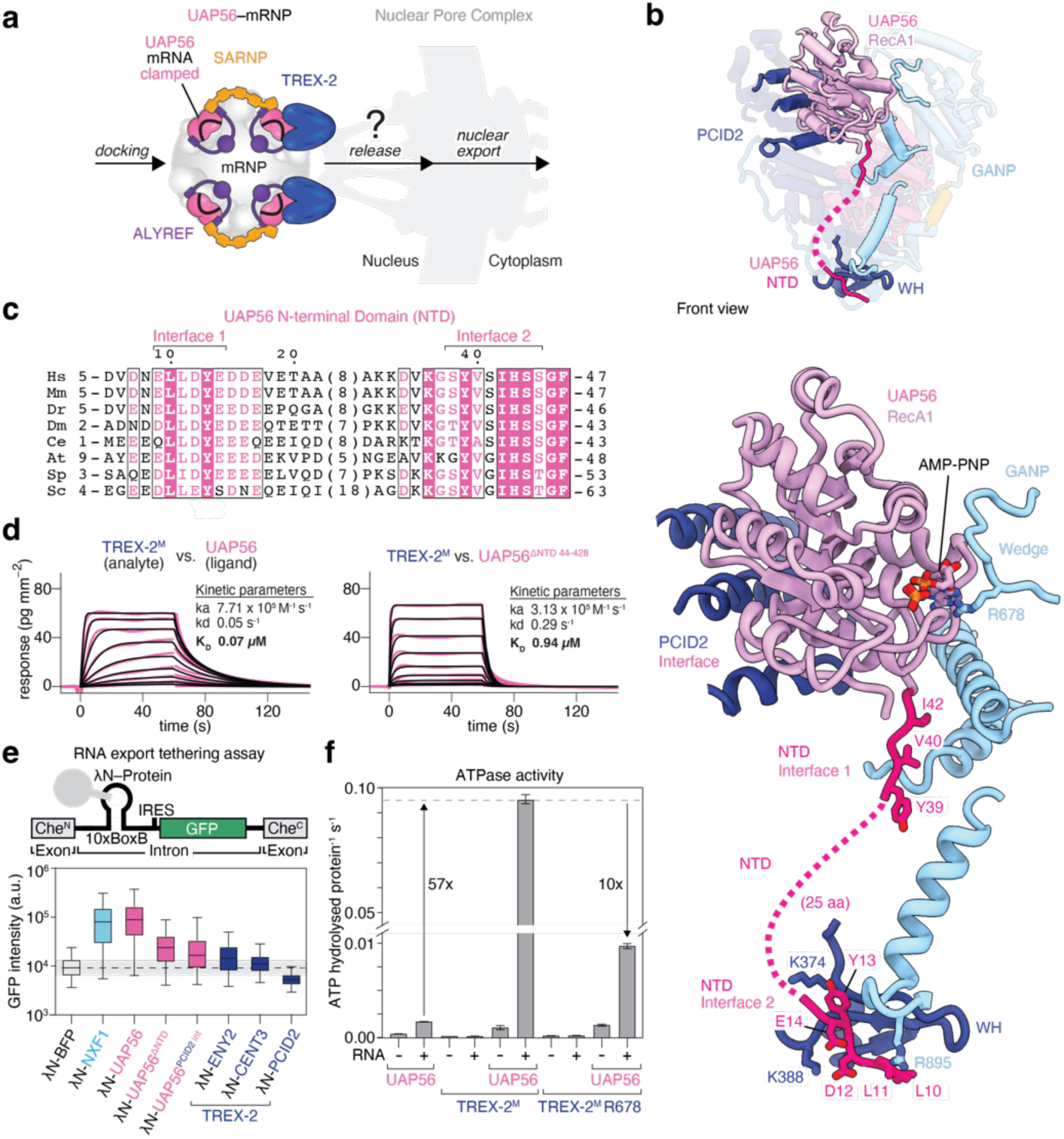
UAP56–TREX-2 interfaces and mRNA release. **a,** Cartoon schematic of an UAP56–mRNP docked at the NPC, which needs to release from TREX-2 through an unknown mechanism to initiate mRNP export. **b,** Details of the UAP56–TREX-2^M^ interfaces. TREX-2 regions not involved in the UAP56 interfaces were omitted for clarity. The inset above shows the UAP56–TREX-2^M^ complex overview. Sidechains of selected interface residues and AMP-PNP are shown as sticks. Colors as in Fig. 3. **c,** Multiple sequence alignment of the UAP56 N-terminal domain (NTD) from *H. sapiens* (Hs, Uniprot ID Q13838), *M. musculus* (Mm, Q9Z1N5), *D. rerio* (Dr, Q803W0), *D. melanogaster* (Dm, Q27268), *C. elegans* (Ce, Q18212*), A. thaliana* (AtRH56, Q9LFN6), *S. pombe* (SpSub2, O13792) and *S. cerevisiae* (ScSub2, Q07478) is shown. Residues invariant or conserved among these species are highlighted in pink or light pink, respectively. **d,** Sensograms for TREX-2^M^ (immobilized) probed with UAP56 or UAP56ΔNTD (residues 44-428). Sensograms (pink line), the fitted model (black), and a binding kinetics summary table are shown. **e,** RNA export tethering assay. The reporter construct is based on an mCherry open reading frame, which was split in two halves (exon 1, exon 2) by an intron containing ten BoxB RNA aptamers, an IRES, and a GFP open reading frame (top). This reporter was stably integrated in a human K562 cell line. λN-tagged proteins are transiently expressed and bind the reporter RNA through the BoxB RNA aptamers. Export of the reporter RNA allows GFP production, which is quantified through Fluorescence-activated cell sorting (FACS) (bottom, see also Supplementary Fig. 10b). Protein levels of transiently expressed λN-UAP56 and λN-UAP56ΔNTD are comparable, shown in Supplementary Fig. 10c. **f,** TREX-2^M^ stimulates UAP56 ATP hydrolysis *in vitro*. Plotted are the ATPase rates (molecules of ATP hydrolyzed protein^-1^ second^-1^) with and without 15 poly-Uridine RNA of UAP56, TREX-2^M^, TREX-2^M^ (R678A), UAP56–TREX-2^M^ or –TREX-2^M^ (R678A) mixtures.

### The conserved N-terminal domain of UAP56 binds TREX-2

UAP56 has an unstructured N-terminal domain (NTD) that is conserved from yeast to humans (Fig. 4c). Although required for mRNA export (Saguez et al., 2013), the molecular function of the NTD is unknown. In the UAP56–TREX-2^M^ structure, the UAP56 NTD binds along the GANP Sac3 domain (via UAP56 residues 39-44, NTD interface II) and between the GANP and PCID2 winged helix (WH) domains (via UAP56 residues 10-15, NTD interface I) (Fig. 4b). Consistent with this, deletion of UAP56 residues 1-28, which removes NTD interface I, resulted in a ∼2.5-fold reduction in UAP56–TREX-2^M^ affinity (K_D_=0.17 µM) compared to full-length UAP56 (K_D_=0.07 µM) (Fig. 4d, Supplementary Fig. 10a). Furthermore, UAP56 lacking the entire NTD (residues 1-43, UAP56ΔNTD) displayed a more than ten-fold reduced affinity (K_D_=0.94 µM) (Fig. 4d, Supplementary Fig. 9c), and an isolated UAP56 NTD peptide (residues 5-23) was sufficient to bind TREX-2^M^ (Supplementary Fig. 9d). Conversely, mutations affecting conserved residues in the WH domains of GANP and PCID2 have been shown to lead to mRNA export defects in yeast *in vivo* (Ellisdon et al., 2012). Contrary to prior speculation, which had attributed these positively charged residues to a putative RNA binding site in TREX-2^M^, our data provide an immediate explanation for these TREX-2^M^ mutations, as they contribute critically to the newly identified interface between TREX-2^M^ and the negatively charged UAP56 NTD (Fig. 4b).

To assess the significance of the identified UAP56–TREX-2 interfaces in a cellular context, we introduced mutations in UAP56 and examined their impact in an RNA export tethering assay. Aptamer-mediated tethering of UAP56 to a reporter pre-mRNA promoted its export to a degree comparable to the tethering of the mRNA export factor NXF1 (Fig. 4e, Supplementary Fig. 10b) (Hargous et al., 2006; Hautbergue et al., 2009; Viphakone et al., 2015), consistent with UAP56 promoting mRNP export after TREX disassembly. The combined mutation of critical residues in the UAP56 NTD interface I (L10S, L11S, D12K, Y13S) resulted in a modest reduction of the export-promoting effect compared to wildtype UAP56. In contrast, the removal of the entire UAP56 NTD strongly diminished its export-promoting capability (Fig. 4e, Supplementary Fig. 10b), in agreement with our corresponding *in vitro* UAP56–TREX-2^M^ affinity measurements (Fig. 4d). This reduction in the export-promoting effect was comparable to mutations of UAP56 RecA1 residues that face PCID2 in our UAP56–TREX-2^M^ structure (D49A, L51W, Q78A, L81K) (Fig. 4e, Supplementary Fig. 10b). Since expression levels and nuclear import of the different λN-tagged UAP56 constructs were unaffected (Supplementary Fig. 10c), we conclude that the observed export defects are due to an impaired UAP56–TREX-2 interaction. Notably, tethering of the TREX-2 subunits PCID2, CETN3, or ENY2 to the reporter pre-mRNA did not promote export (Fig. 4e), presumably because human TREX-2 is a stable complex that is constitutively anchored to the NPC basket (Umlauf et al., 2013).

Collectively, these findings suggest that the transient association of UAP56 with the NPC-anchored TREX-2 complex stimulates nuclear mRNA export. *In vivo*, efficient docking at the NPC may be further enhanced by multivalent interactions between multiple UAP56 molecules of the mRNP and multiple TREX-2 complexes, which may be bound to the NPC in up to eight copies owing to the 8-fold symmetry of the NPC.

### TREX-2 triggers RNA release from UAP56

To allow export through the NPC, mRNPs must eventually dissociate from TREX-2, which is anchored to the NPC basket (Fig. 4a). An important clue as to how this might happen came from our UAP56–TREX-2^M^ structure. Although we prepared the UAP56–TREX-2^M^ cryo-EM sample with a UAP56–UCM–N-UBM fusion protein and in the presence of RNA and non-hydrolysable AMP-PNP, UAP56 is not clamped on RNA in the structure (Figs. 3e, 4b). Instead, the UAP56 RecA1 lobe is sandwiched between PCID2 and a highly conserved loop within GANP (residues 674-686) (Figs. 3e, 4b; Supplementary Fig. 9b,e). The GANP loop, which we named the ‘wedge’, is only visible in the new UAP56–TREX-2^M^ complex structure, but not in the isolated human TREX-2^M^ (Fig. 3e), or in a published yeast TREX-2^M^ cryo-EM structure (Gordon et al., 2017) (Supplementary Fig. 9a). In the UAP56–TREX-2^M^ complex, the GANP wedge occupies the position near the UAP56 RecA1 lobe, which is normally occupied by the RecA2 lobe in RNA-clamped UAP56. Notably, the UAP56–TREX-2^M^ complex does contain the AMP-PNP nucleotide, which is bound between UAP56 RecA1 residue F65 and the evolutionarily invariant GANP wedge residue R678 (Fig. 4b; Supplementary Fig. 9e,f). In our structure, GANP R678 mimics F381 of UAP56 RecA2, which coordinates the nucleotide in RNA-clamped UAP56 (Fig. 4b, Supplementary Fig. 9f) (Ren et al., 2017; Xie et al., 2023). These data suggest a compelling model in which the GANP wedge promotes the release of mRNA from UAP56, explaining previous observations implicating TREX-2 in the removal of UAP56 from mRNPs in yeast (Wilmes et al., 2008).

Since the release of RNA from DExD-box helicases is typically coupled to ATP hydrolysis and/or ADP and P_i_ release, we determined whether TREX-2^M^ affects the ATPase activity of UAP56. Using an *in vitro* ATPase assay (Fig. 4f, Supplementary Fig. 10d), we observed that wild-type TREX-2^M^ stimulates the ATPase activity of UAP56 by more than fifty-fold in the presence of RNA (Fig. 4f, Supplementary Fig. 10d). A single point mutation of the GANP wedge residue R678 to alanine reduced the ATPase stimulatory effect by approximately ten-fold (Fig. 4f, Supplementary Fig. 10d), without affecting the binding of UAP56 to mutant TREX-2^M^ (Supplementary Fig. 9c). Taken together, these data suggest that TREX-2 serves not only as the docking site for UAP56–mRNPs at the NPC, but also as the site where UAP56 dissociates from mRNPs.

## DISCUSSION

### A general model for mRNA export

The data presented in this study offer a comprehensive framework for understanding mRNA export, revealing a general and conserved process (Fig. 5). Central to this model is the ATPase UAP56, which orchestrates a linear process that guides mRNAs through distinct molecular complexes, from the completion of mRNP biogenesis to the loading of the mRNA export factor NXF1–NXT1 at the NPC. Synthesizing previous insights, our results delineate a five-step pathway that elucidates the sequential events governing mRNA export (I-V, Fig. 5).

**Figure 5.**
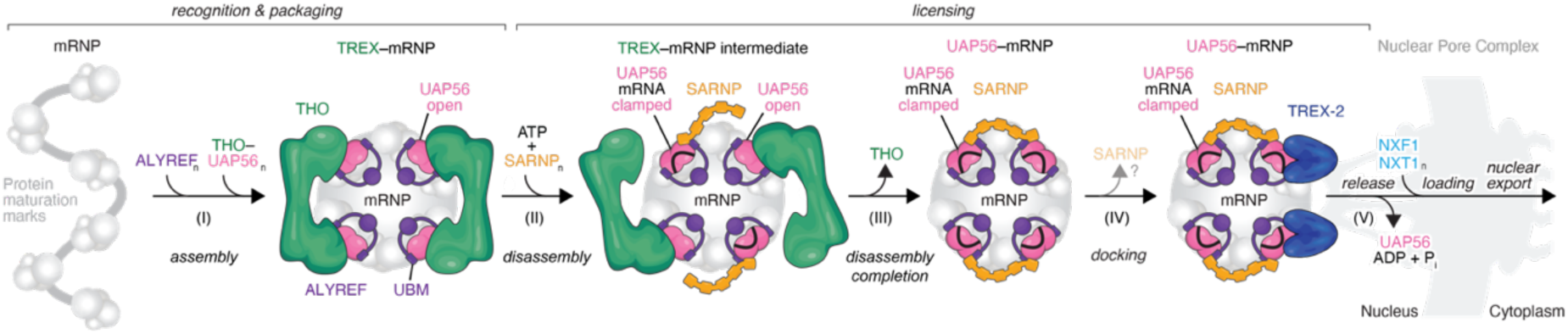
Model for licensing mRNA export. The ATPase and mRNA clampase UAP56 is a central molecular switch that steers human mRNAs through mRNP packaging and multivalent TREX–mRNP globule assembly (I), TREX–mRNP disassembly and the formation of export-competent UAP56–mRNPs, assisted by the multivalent protein SARNP (II and III), mRNP docking (IV), and mRNP release (IV) at the NPC via TREX-2. Loading of the mRNA export factor NXF1–NXT1 onto mRNPs may occur in the nucleoplasm (Schmidt et al., 2009) or at the NPC (Ben-Yishay et al., 2019; Derrer et al., 2019), initiating mRNP nuclear export.

First, during mRNA maturation, ALYREF and other mRNA export adaptors (Ohno et al., 2000; Richardson et al., 2004; Hautbergue et al., 2009; Chang et al., 2013; Viphakone et al., 2015; Gromadzka et al., 2016) bind the newly made mRNP via specific marks, which initiates the selective packaging of mRNA into mRNP globules through low affinity and multivalent protein-protein and protein-mRNA interactions (Köhler and Hurt, 2007; Singh et al., 2015; Metkar et al., 2018; Pacheco-Fiallos et al., 2023; Bonneau et al., 2023).

Second, these mRNPs acquire a high density of N-UBMs and C-UBMs on their surface, which recruit the tetrameric THO complex via the four bound UAP56 molecules, thus assembling TREX on the mRNP surface (Pacheco-Fiallos et al., 2023). TREX thereby aids further mRNP compaction and chaperones the mRNA, preventing the formation of harmful R-loops.

Third, THO releases from these multivalent TREX–mRNPs when UAP56 clamps onto mRNA together with ATP. The pentavalent protein SARNP may bind cooperatively with ALYREF in the resultant UAP56–mRNP complexes to simultaneously prevent UAP56 from reassociating with THO and thereby assist TREX disassembly.

Fourth, these remodeled UAP56–mRNPs then diffuse in the nucleus (Grünwald and Singer, 2010; Mor et al., 2010) before docking at the NPC-anchored TREX-2 complex via UAP56. Once docked, UAP56–mRNPs could bind the mRNA export factor, NXF1–NXT1 that is enriched at the NPC by several FG repeat-containing proteins (Katahira et al., 1999; Sträßer et al., 2000; Grant et al., 2002; Weis, 2002) that include the TREX-2 subunit GANP (Fischer et al., 2002; Dimitrova et al., 2015).

Fifth, ATP hydrolysis and TREX-2 unclamp UAP56 from mRNPs, releasing the mRNPs directly at the nuclear envelope for export through the NPC via NXF1–NXT1. Consistent with this model, overexpression of the isolated GANP Sac3 domain in yeast leads to an mRNA export defect (Fischer et al., 2002), presumably because nucleoplasmic TREX-2 complexes prematurely release UAP56 from mRNPs.

This general mRNA export model relies on UAP56 as the pivotal molecule and a series of multivalent protein-protein and protein-mRNA interactions that provide numerous opportunities for the regulation of mRNA export. The *in silico* UAP56 protein interaction screen uncovered novel UBM-containing and UCM-containing proteins, including a protein of viral origin, as well as TREX-2 complex paralogs (Supplementary Fig. 6i,j, Supplementary Table 4). These results highlight the potential for regulation at each step of the export pathway, and at the same time demonstrate the power of *in silico* protein-protein interaction screens to generate new hypotheses.

Forty years ago, Günter Blobel proposed the ‘gene gating’ hypothesis, suggesting that actively transcribed genes would localize to the NPC via a specific DNA element, enhancing gene expression efficiency (Blobel, 1985). Subsequent research demonstrated that the NPC-anchored TREX-2 complex participates in gene gating processes in yeast and humans (Fischer et al., 2004; Rodríguez-Navarro et al., 2004; Cabal et al., 2006; Kurshakova et al., 2007). However, no responsible DNA element was identified. Instead, several DNA-associated co-activator complexes, including SAGA and Mediator, were implicated (Rodríguez-Navarro et al., 2004; Cabal et al., 2006; Kurshakova et al., 2007; Schneider et al., 2015). Our data suggest an alternative model. The interaction between UAP56 and TREX-2 offers insight into how highly transcribed genes might associate with the NPC, not through DNA elements, but through features of the mRNA. During transcription, mRNA packaging and TREX disassembly could potentially load UAP56 onto nascent mRNAs (Sträßer and Hurt, 2001; Zenklusen et al., 2002). Consequently, actively transcribed genes may harbor DNA-tethered UAP56–mRNPs, which collectively enrich at the NPC through TREX-2, facilitating efficient mRNA nuclear export. Supporting this model, mutations in yeast GANP that affect gene gating (Schneider et al., 2015), map to the UAP56–TREX-2 interface in our obtained structure (Supplementary Fig. 9b,e).

In conclusion, our combined results reveal a mechanistic framework for the selective and efficient export of mRNA and elucidate the molecular functions of conserved proteins and multi-subunit complexes that control individual steps. At the core of this pathway lies the protein UAP56, which orchestrates the nuclear export of mRNA through an ATP-gated molecular switch.

## Acknowledgments

We thank our colleagues in the Brennecke and Plaschka groups for their support and discussions; staff at the Protein Technologies Facility at the Vienna BioCenter Core Facilities (VBCF), a member of the Vienna BioCenter (VBC), for assistance with protein production; staff at the VBCF Electron Microscopy Facility, in particular T. Heuser and H. Kotisch, for support, data collection and maintaining facilities; V.-V. Hodirnau at the Institute of Science and Technology Austria EM facility for cryo-EM data collection; staff at the VBCF Proteomics facility, in particular E. Roitinger and O. Hudecz, for mass spectrometry analysis; R. Zimmermann and his team for computational support; M. Madalinski for peptide synthesis; staff at the in-house Molecular Biology Service for reagents; S. Ameres and J. Zuber for sharing reagents; S. Ameres and U. Elling for help with export reporter design; A. Phillip for help with mammalian cell culture; M. Vorländer and R. Faraway for help with data analysis; C. Bernecky, L. Lorenzo-Orts, and I. Patten for discussions; and A. Anderson (Life Science Editors), C. Bernecky, T. Heick-Jensen, L. Lorenzo-Orts, A. Stark and members of the Brennecke and Plaschka groups for feedback on the manuscript. U.H. was supported by a Marie Sklodowska-Curie fellowship (896416) and an EMBO long-term fellowship (ALTF_1175-2019). D.R.-B. was supported by a Marie Sklodowska-Curie fellowship (101028744). J.B. by the Austrian Academy of Sciences and Austrian Science Fund (W1207). Research in the laboratory of C.P. is supported by Boehringer Ingelheim, the European Research Council under the Horizon 2020 research and innovation programme (ERC-2020-STG 949081 RNApaxport) and by the Austrian Science Fund (FWF) doc.funds program DOC177-B (RNA@core: Molecular mechanisms in RNA biology). The funders had no role in study design, data collection and analysis, decision to publish or preparation of the manuscript. For the purpose of open access, the author has applied for a CC BY public copyright licence to any Author Accepted Manuscript version arising from this submission.

## Author contributions

U.H. designed research and with M.G., L.F., and T.P. purified recombinant proteins. U.H. and M.G. performed biochemical experiments and collected TREX-2 cryo-EM data. B.P.-F. performed the native TREX disassembly assay and collected TREX–EJC–ALYREF cryo-EM data. U.H. and C.P. carried out cryo-EM data analysis and structure determination. U.S. performed the export tethering assay, for which M.-F.S. engineered and M.-F.S. and C.P. designed the cell line. D.R.-B and L.T. engineered and grew mammalian cell lines. U.H. performed structural predictions with a pipeline implemented by D.H.. U.H., J.B., and C.P. prepared the manuscript with input from all authors. J.B. supervised L.T. and D.H. and designed research. J.B. and C.P. supervised U.H.. C.P. supervised M.G., U.S., B.P-F., L.F., D.R-B., T.P., M.-F. S., designed research, and initiated the mRNA export project.

## Data availability

Three-dimensional cryo-EM density maps of the TREX–EJC–ALYREF complex, TREX-2M and UAP56–TREX-2M have been deposited into the Electron Microscopy Data Bank under the accession numbers EMD-18980 (Map-A) and EMD-18979 (Map-B), EMD-18977 (Map-C), EMD-18978 (Map-D) and EMD-18981 (Map-E) respectively. The coordinate files of the TREX– EJC–ALYREF, TREX-2M and UAP56–TREX-2M have been deposited into the Protein Data Bank under the accession numbers 8R7L, 8R7J, and 8R7K. The coordinate file of the TREX– mRNA complex was updated in the Protein Data Bank under the accession number 7ZNK.

## MATERIALS AND METHODS

### Vectors and sequences

All vectors and sequences are described in Supplementary Table 5.

### Protein purification

#### THO complex and EJC

Recombinant THO complex tetramer (THOC1, THOC2 residues 1–1203, THOC3, THOC5, THOC6 and THOC7), dimer (same as tetramer but lacking THOC6) and monomer (THOC1, THOC2 residues 1–1203, THOC3, THOC5 residues 1–224, THOC7 residues 1–159) as well as the EJC subunits eIF4A3, MAGOH–Y14, ALYREF^N^ (residues 1-183 were purified as described previously (Pühringer et al., 2020; Pacheco-Fiallos et al., 2023).

#### UAP56 and UAP56 fusion proteins

6xHis-TwinSTREPII-3C-UAP56, 6xHis-MBP-3C-UAP56, 6xHis-3C-UAP56ΔNTD (residues 44-428) and 10xHis-3C-UAP56 D283R constructs were purified as described previously for UAP56 (Pühringer et al., 2020). The fusion proteins 10xHis-UAP56–UCM-1, 10xHis-UAP56–N-UBM, 10xHis-UAP56–UCM-1–N-UBM were expressed in E. coli BL21 DE3 RIL cells grown in LB medium, induced at OD 600nm of 1.0 with 0.5 mM IPTG and incubated at 37°C for 3h. Cells were resuspended in lysis buffer (25 mM HEPES pH 7.9, 5% (v/v) glycerol, 300 mM NaCl, 20 mM imidazole, 0.05% Tween20 and cOmplete EDTA-free protease inhibitor cocktail) and lysed by sonication. The lysate was clarified by centrifugation and the supernatant was filtered through 1µm and 0.45 µm filters and applied to a HisTrap HP 5 mL column (Cytiva), pre-equilibrated with buffer A (25 mM HEPES pH 7.9, 5% (v/v) glycerol, 300 mM NaCl, 20 mM imidazole). The column was washed with buffer A containing 44 mM imidazole and proteins were eluted with a linear gradient from 50 mM to 300 mM imidazole. Peak fractions were diluted with buffer C (25 mM HEPES pH 7.9, 5% (v/v) glycerol, 1 mM DTT) to 100 mM NaCl and further purified by anion-exchange chromatography using a HiTrapQ 5 mL column (Cytiva), pre-equilibrated in buffer C. The column was washed with buffer C containing 100 mM NaCl and eluted using a linear gradient from 200 mM to 400 mM of NaCl. Peak fractions were concentrated and loaded on a HiLoad 16/600 Superdex 200 pg column (Cytiva) equilibrated using buffer D (25 mM HEPES pH 7.9, 5% (v/v) glycerol, 250 mM NaCl, 1 mM DTT). The purified proteins were concentrated, flash frozen and stored at −80°C.

#### SARNP UCM-1 and ALYREF N-UBM

10xHis-SUMO-3V5 tagged ALYREF N-UBM, SARNP UCM-1, UCM-1–N-UBM and UCM-1(R106D)–N-UBM were expressed in *E. coli* BL21 DE3 RIL cells. UCM-1, N-UBM and UCM-1 (R106D) were expressed in LB media at 37°C for 3 h after induction with 0.5 mM IPTG at OD 600 nm of 1.0. UCM-1–N-UBM and UCM-1(R106D) –N-UBM were incubated at 18°C overnight (o/n) upon induction. Cell pellets were resuspended in lysis buffer and lysed by sonication. Lysates were clarified by centrifugation, filtered through 1 µm and 0.45 µm filters and loaded on a HisTrap HP 5 mL column equilibrated in buffer A. The column was washed with buffer A and proteins were eluted at 350 mM imidazole. Peak fractions were diluted to 50 mM NaCl with buffer C and loaded onto a HiTrapQ HP 5 mL column equilibrated in buffer C. The column was washed with buffer C supplemented with 50 mM NaCl and eluted using a linear gradient from 50 mM to 500 mM NaCl. Peak fractions were concentrated and applied to a HiLoad 16-600 Superdex 75 pg column (Cytiva) equilibrated using buffer E (10 mM HEPES pH 7.9, 500 mM NaCl, 10% (v/v) glycerol, 1 mM DTT, 20 mM imidazole). The peak fractions were concentrated again, flash frozen in liquid nitrogen and stored at −80°C. Buffer A and B for the purification of UCM-1–N-UBM contained 500 mM NaCl.

The UCM-1 R106D mutant was purified using a similar strategy with the following exceptions: two wash steps were performed during HisTrap using buffer A including a high salt wash (25 mM HEPES pH 7.9, 5% (v/v) glycerol, 1 M NaCl) and buffer A supplemented with 50 mM imidazole. During the anion-exchange step, the column was washed with 100 mM NaCl and eluted by a linear gradient from 100 mM to 400 mM NaCl.

#### SARNP

SARNP-6xHis was expressed in *E. coli* BL21 DE3 RIL cells grown in LB medium o/n at 18°C after induction with 0.5 mM IPTG at OD_600_=1.0. Cell pellets were resuspended in lysis buffer (50 mM HEPES pH 7.9, 500 mM NaCl, 10% (v/v) glycerol, 20 mM imidazole, 1 mM DTT, 0.5 mM PMSF, cOmplete EDTA-free protease inhibitor cocktail and 0.1% Tween20), lysed by sonication and centrifuged. The supernatant was filtered through a 0.4 µm filter and loaded on a HisTrap HP 5 mL column equilibrated using buffer E. The column was washed and SARNP was eluted using two linear gradients from 0 to 100 mM and 100 to 500 mM imidazole. Peak fractions were diluted to 200 mM NaCl using buffer F (25 mM HEPES pH 7.9, 10% (v/v) glycerol and 2.5 mM DTT) and applied to a HiTrapQ HP 5 mL column equilibrated using buffer F (200 mM NaCl). The column was washed and eluted over a linear gradient from 200 to 700 mM NaCl. Peak fractions were concentrated and applied to a HiLoad 16-600 Superdex 200 pg column, pre-equilibrated using buffer D containing 2.5 mM DTT and 250 mM salt. The purified protein was concentrated, flash frozen and stored at −80°C.

#### TREX-2^M^ and TREX-2^M^ (R678A)

TREX-2^M^ and TREX-2^M^ (R678A) were expressed in *E. coli* BL21 DE3 RIL cells grown in LB medium at 37°C until OD 600 nm at 1.0. Expression was induced by addition of 0.5 mM IPTG and cells were incubated at 18°C overnight. Cells were harvested by centrifugation and resuspended in lysis buffer (containing 500 mM NaCl and no Tween20 for the TREX-2^M^ purification). Cells were lysed by sonication and lysates were centrifuged. The supernatant was filtered through 1 μm and 0.45 μm filters and applied to a HisTrap HP 5 mL column equilibrated with buffer A, washed with buffer A (50 mM NaCl) and eluted over a linear gradient to 350 mM imidazole. The complex was diluted in buffer C to 100 mM NaCl and loaded on a HiTrapQ HP 5 mL column equilibrated with buffer C containing 100 mM NaCl. Following a wash step with the buffer C containing 100 mM NaCl, the complex was eluted from the HiTrapQ column using a linear gradient to 800 mM NaCl (500 mM NaCl for TREX-2^M^ (R678A)). Peak fractions were concentrated and applied to a HiLoad 16-600 Superdex 200 pg column equilibrated with buffer D. The purified complex was concentrated, flash frozen and stored at −80°C.

#### MBP-GANP and PCID2–UAP56–UCM-1–N-UBM

MBP-GANP (residues 582-1004) and 10xHis-PCID2–UAP56–UCM–N-UBM – SEM1 were expressed in *E. coli* BL21 DE RIL cells. MBP-GANP was expressed in LB media at 18°C o/n upon induction with 0.5 mM IPTG at OD 600 nm at 1.0 and 10xHis-PCID2–UAP56–UCM-1–N-UBM – SEM1 was expressed in autoinduction medium at 30°C. Bacterial cell pellets for MBP-GANP were lysed in buffer A containing 500 mM NaCl by sonication and centrifuged. The supernatant was loaded on a HisTrap HP 5 mL column equilibrated using buffer A containing 500 mM NaCl and 50 mM imidazole. The column was washed with this buffer A and eluted over a linear gradient to 350 mM imidazole using buffer B contained 500 mM NaCl. Peak fractions were diluted to 100 mM NaCl using buffer C, applied to a HiTrapQ HP 5 mL column, and washed with buffer C containing 100 mM NaCl. The proteins were eluted using a linear gradient to 800 mM NaCl. The flow-through of the anion-exchange step was concentrated and loaded on a HiLoad 16-600 Superdex 200 pg column equilibrated with buffer E. Peak fractions were concentrated, flash frozen and stored at −80°C.

10xHis-PCID2–UAP56–UCM-1–N-UBM – SEM1 was purified using a similar strategy with the following changes: Buffers A and B contained 300 mM NaCl and the co-expressed complex was eluted from HisTrap using a linear gradient from 50 to 300 mM imidazole. Moreover, the column was washed with buffer C containing 160 mM NaCl during the anion-exchange and eluted over a linear gradient from 160 to 400 mM NaCl.

#### MBP-MCP

MBP-MCP was expressed in *E. coli* Rosetta2 pLysS cells, grown in LB medium at 37°C until OD 600 nm at 0.7, induced by addition of 0.5 mM IPTG and incubated at 37°C for 3h. Cells were resuspended in lysis buffer (20 mM HEPES pH 7.9, 200 mM KCl, 1 mM EDTA, 0.5 mM PMSF), lysed by sonication. The lysate was centrifuged, filtered through a 0.45 µm filter and loaded on a MBP Trap HP column (Cytiva) equilibrated with buffer G (20 mM HEPES pH 7.9, 200 mM KCl, 1 mM EDTA). The column was washed first with buffer G and then with buffer H (20 mM HEPES pH 7.9, 20 mM KCl, 1 mM EDTA) and the protein was eluted using buffer H containing 10 mM maltose. The protein was further purified using a HiTrap Heparin HP 5 mL column and washed with buffer H (no EDTA). The protein was eluted over a linear gradient to 400 mM KCl. Peak fractions were flash frozen in storage buffer (10 mM HEPES pH 7.9, 57 mM KCl, 1 mM EDTA, 10 % (v/v) glycerol) and stored at −80°C.

### Pull down experiments using recombinant proteins

#### In vitro THO–UAP56 disassembly assay

Recombinant MBP-UAP56 (6.75 μg per reaction) was combined with a two-fold molar excess of monomeric THO complex (10 μg per reaction) in buffer I (20 mM HEPES pH 7.9, 50 mM KCl, 1 mM MgCl_2_, 5 % glycerol, 0.1% Igepal CA-630) and incubated with 30 μl of amylose resin (E8021S, NEB), pre-equilibrated in buffer I, for 30 min at room temperature (RT). The resin was then separated from the supernatant by centrifugation, washed three times with buffer I, resuspended in 40 μl of buffer I per reaction, and split into individual tubes for each THO– UAP56 disassembly reaction. Components for the release reaction were prepared in a final volume of 40 μl buffer I (final concentrations / amounts: 200 μM 15 U RNA, 0.1 mM ATP, 55/55/60 μg Sumo-V5–N-UBM / Sumo-V5–UCM-1 / Sumo-V5–UCM-1–N-UBM), combined with the amylose resin with immobilized UAP56–THO complex and incubated for 60 min at RT. After four washes with buffer I the bead-retained complexes were then eluted in buffer I supplemented with 100 mM maltose for 20 min at RT. Elutions and input samples of the individual recombinant proteins were separated on 4 – 12% gradient SDS-PAGE gels and visualized by Coomassie staining. The amount of bead-retained THO complex in each reaction was analyzed in Fiji (Schindelin et al., 2012). The intensity of the THOC2 band was measured, normalized to the intensity of the MBP-UAP56 band, and the background subtracted; THOC2 in the reaction incubated with buffer I without supplements was set to 100 %.

#### RNA clamping assay

In step 1, for each reaction 1 μg of in vitro transcribed 450 nt AdML RNA and 5.1 μg of MBP- MS2 (equimolar with the RNA) in buffer J (20 mM HEPES pH 7.9, 100 mM KCl, 2 mM MgCl_2_, 5 % Glycerol, 0.1% Igepal CA-630) were incubated with 20 μl of amylose resin (E8021S, NEB), pre-equilibrated in buffer J for 30 min at RT. The resin was then harvested by centrifugation, the supernatant containing unbound components removed, and three washes with buffer J conducted before the resin with immobilized MBP-MS2–RNA was resuspended in 40 μl buffer J supplemented with 1 mM AMP-PNP and split into the desired number of reactions. Components to be tested for RNA binding in step 2 (23 μg UAP56 or UAP56–N-UBM, UAP56–UCM, UAP56–UCM-1–N-UBM {2-fold molar excess over the RNA}, 24 μg SARNP {2-fold molar excess over UAP56}) were prepared in buffer J containing 1 mM AMP-PNP, combined with the resin prepared in step 1 and incubated for 90 min at RT. The resin was then again harvested by centrifugation, washed three times with buffer J and incubated with 40 μl buffer J containing 0.4 μg benzonase to elute RNA-bound proteins. Elutions and input samples of the individual recombinant proteins were separated on 4 – 12% gradient SDS-PAGE gels and visualized by Coomassie staining. To assess the amount of RNA-bound UAP56 in Fiji (Schindelin et al., 2012) we measured the intensity of the UAP56 band, subtracted the background and normalized to UAP56 in the presence of SARNP set to 100 %.

#### SARNP UCM-1 and ALYREF N-UBM - UAP56 pulldown

To assess the interaction of UAP56 and the SARNP UCM-1 or the ALYREF N-UBM 7.5 μg of Sumo-V5-3C tagged UCM-1, N-UBM or UCM-1–N-UBM were combined with a 4-fold molar excess of UAP56 in buffer K (25 mM HEPES pH 7.9, 40 mM NaCl, 5% glycerol, 0.01% Igepal CA630, 1 mM MgCl_2_, 1 mM TCEP) in the presence of 50 μM 15 U RNA and 1 mM AMP-PNP, and incubated for 1h at 4 °C before being added to 10 µl magnetic V5 beads (v5tma, Chromotek), pre-equilibrated in buffer K. After incubation for another hour rotating at 4 °C the beads were centrifuged briefly to recover beads from the lid (1300 x g, 2 min, 4°C) and washed three times with buffer K on a magnetic rack. Samples were eluted using 30 μL 200 mM glycine (pH 2.52) for 5 min at RT. Eluates were neutralized using 2.5 μL 1 M Tris pH 10.4, and, together with input samples of the individual recombinant proteins, separated on 4 – 12% gradient SDS-PAGE gels and visualized by Coomassie staining.

#### UAP56 - TREX-2^M^ pulldown

To analyze the interaction of TREX-2^M^ and UAP56, TREX-2^M^ with an MBP-tag on the GANP subunit, was combined with a 4-fold molar excess of UAP56 and a 10-fold molar excess of UCM–UBM fusion peptide in buffer K, with or without 50 μM 15U RNA and 1 mM AMP-PNP, and incubated rotating for 1 h at 4 °C. Samples were added to 30 μl pre-equilibrated amylose resin (E8021S, NEB), and incubated for another hour rotating at 4 °C. Beads were centrifuged (1300 x g, 2 min, 4°C) to remove the unbound fraction, washed three times with buffer K and bead-bound complexes eluted for 1h at RT in 30 μl buffer K supplemented with 100 mM maltose. Elutions and input samples of the individual recombinant proteins were separated on 4 – 12% gradient SDS-PAGE gels and visualized by Coomassie staining.

#### UAP56 NTD –TREX-2^M^ pulldown

To test the interaction of the isolated UAP56 NTD and TREX-2^M^, 30μl of Pierce™ High Capacity NeutrAvidin™ Agarose beads (# 29202, ThermoScientific) were pre-equilibrated with buffer K and incubated with or without 30 μg of biotinylated UAP56 NTD peptide in buffer K for 1h at room temperature. The beads were then washed three times to remove unbound peptide and incubated with protein samples (set up in a 50 µl reaction containing 50 µM 15U RNA and 1 mM AMP-PNP and, as applicable: 7.5µg TREX-2^M^ +/- a 4-fold molar excess of UAP56; 7.5µg GANP(582-1004) with a 2.5-fold molar excess of the PCID2-UAP56-UCM-N-UBM–SEM1).

After an incubation of 1 h rotating at 4 °C, the beads were again collected by centrifugation, washed three times with buffer K and bead-bound material eluted for 5 min at RT in 30 μl of 200 mM glycine pH 2.52. The elutions were neutralized with 100 mM Tris pH 10.4, separated alongside input samples of isolated recombinant proteins on 4 – 12% gradient SDS-PAGE gels and visualized by Coomassie staining.

### Grating Coupled Interferometry (GCI)

GCI experiments were performed on a Creoptix WAVE system (Creoptix AG, Switzerland) using 4PCP WAVEchips (quasi-planar polycarboxylate surface; Creoptix AG, Switzerland). Chips were conditioned with borate buffer (100 mM sodium borate pH 9.0, 1 M NaCl), and either strepdavidin (10 μg ml^-1^ in 10 mM sodium acetate pH 5.0) or a monoclonal anti-V5 antibody (R960252, Invitrogen; 2 μg ml^-1^ in 10 mM sodium acetate pH 5.0) immobilized using a standard amine coupling protocol, followed by passivation of the surface with BSA (0.5 % in 10 mM sodium acetate pH 5.0) and final quenching with 1 M ethanolamine pH 8.0. Biotinylated 15 U RNA, UCM or UBM peptides or V5-tagged THO or TREX-2^M^ complexes were captured on the prepared chip until the desired density was reached. UAP56 was injected in a 1:2 dilution series, starting from a highest concentration of 5 μM with or without 200 μM 15 U RNA, in 25 mM HEPES pH 7.9, 50 mM KCl, 1 mM MgCl_2_, 1 mM TCEP, with and without 1 mM ATP at 25°C.

Blank injections were used for double referencing and a DMSO calibration curve was used for bulk correction. Analysis and correction were performed using the Creoptix WAVEcontrol software (applied corrections: X and Y offset, DMSO calibration, double referencing) using a one-to-one binding model. The data and fitted models were plotted in R.

### GFP-UAP56 immunoprecipitation

GFP immunoprecipitations were performed in triplicates from nuclei of GFP-UAP56 or wildtype K562 cells. For each replicate 200 mio cells were fractionated into nuclei and cytoplasm as previously described (Suzuki et al., 2010; Pacheco-Fiallos et al., 2023), nuclei lysed in buffer L (50 mM Tris pH 7.5, 100 mM KCl, 3 mM MgCl_2_, 0.25% Triton X-100, 0.25% Igepal CA630, 10% glycerol, 1x protease inhibitor cocktail, 1mM DTT) supplemented with 1 ug ml^-1^ benzonase and 0.1 % deoxycholate, and the lysate incubated for 15 min at 4 °C on a rotating wheel followed by a centrifugation step to pellet chromatin (21,000 x g, 10 min, 4 °C). The supernatant was then incubated with 20 μl magnetic GFP-Trap MA-Agarose beads (Chromotek), pre-equilibrated in buffer L, and incubated on a rotating wheel for 4 h at 4 °C. The beads were then collected on a magnetic rack, washed four times in 1 ml buffer L, and four times with 20 mM Tris pH 7.5,

100 mM KCl. After the final wash step all buffer was removed and the beads snap-frozen in liquid nitrogen. The samples were analyzed by mass spectrometry starting from an on-bead digest of bound protein complexes.

### Endogenous TREX-disassembly assay

Nuclear Extract (NE) from a THOC1-3C-GFP overexpressing K562 cell line was prepared as previously described (Pühringer et al., 2020). 3.6 ml of NE were supplemented with protease inhibitor cocktail and incubated with GFP-Trap Agarose resin (Chromotek), pre-equilibrated with buffer M (20 mM HEPES pH 7.9, 100 mM KCl, 2 mM MgCl2, 8 % glycerol, 0.05 % (v/v) Igepal CA-630, 0.5 mM TCEP), for 3 h at 4 °C. The beads with immobilized endogenous TREX-mRNPs were then washed five times with 1.5 ml buffer M, aliquoted in twelve individual reactions and harvested by centrifugation for 1 min at 1000xg to remove the supernatant. Meanwhile, 10xHis-Sumo-3V5-3C-UBM, 10xHis-Sumo-3V5-3C-UCM or 10xHis-Sumo-3V5- 3C-UCM–N-UBM were prepared at a final concentration of 19 µM in buffer G (25 mM HEPES pH 7.9, 200 mM NaCl, 10 mM MgCl2, 10 % glycerol, 5 mM ATP, 1 mM TCEP) and incubated at room temperature for 30 min. Next, the beads with immobilized endogenous TREX-mRNPs were incubated either with buffer G or supplemented with UCM and / or UBM peptide in buffer G as described for 1 h at room temperature with rotation. After addition of a final concentration of 50 µg/ml benzonase and a further 30 min incubation, the beads were centrifuged for 1 min at 1000xg and washed twice with buffer M. Complexes remaining on the beads were eluted by boiling in 2x SDS sample buffer, loaded in an SDS–PAGEs and ran for 3min at 180V in 1XMOPS Buffer. The gels were stained with Coomassie blue, and the bands containing bead-retained protein were excised for Mass Spectrometry analysis.

An aliquot of the elutions was analyzed by western blotting following standard protocols. We used anti-GFP (CAS A11122, ThermoFisher, diluted 1:1000), anti-UAP56 (AB181059, Abcam diluted 1:1000), and anti-EIF4A3 (AB180519, Abcam, diluted 1:1000). Primary antibodies were incubated overnight at 4 °C. For detection, we used a secondary goat-anti-rabbit antibody coupled to HRP (CAS 31466) and developed the membrane with Pierce ECL reagent (CAS 32106).

### CryoEM sample preparation, imaging, and analysis

#### Model building for the endogenous human TREX complex including the ALYREF N-UBM

The structure of the human endogenous TREX complex (PDB ID 7ZNK) (Pacheco-Fiallos et al., 2023) was analyzed together with the THO monomer 2B map (EMDB-14806) (Pacheco-Fiallos et al., 2023). Manual inspection revealed additional density on the UAP56 RecA2 lobe, which we hypothesized to be the ALYREF N-UBM. The ALYREF N-UBM was modelled in Coot based on the superposition of an AlphaFold2 Multimer prediction model of a UAP56–ALYREF complex on UAP56 chain p. All structural figures were prepared using UCSF Chimera X (Goddard et al., 2018; Pettersen et al., 2021).

#### TREX–EJC–RNA complex reconstitution and sample preparation

TREX–EJC–RNA complexes were reconstituted as described previously (Pacheco-Fiallos et al., 2023) with small modifications. We used a 15 U ssRNA to assemble the ALYREF^N^–EJC–RNA and ATPγS was omitted from Buffer U. The eluted sample was loaded on a 15-40% sucrose density gradient and centrifuged at 23,000rpm for 16h in a SW60Ti rotor. We collected fractions and analyzed every other fraction by SDS–PAGE stained with Coomassie blue.

For Cryo-EM sample preparation, we followed the described methodology in ref. (Pacheco-Fiallos et al., 2023) (Methods, *Sample preparation of TREX–EJC–RNA*) with the following variations: the 15-40% sucrose density gradient was supplemented with a glutaraldehyde gradient from 0 to 0.05% to stabilize the complexes and it was centrifuged at 23,000rpm for 16h in a SW60Ti rotor, and we applied the sample to glow discharged Quantifoil Cu 200 2/1 grids.

#### Cryo-EM data acquisition of TREX–EJC–RNA complex reconstituted on 15 U RNA

Data were collected at IST Austria on a Thermo Fisher Titan Krios G3i operated at 300 keV, equipped with a Gatan K3 direct electron detected operated in counting mode and a BioQuantum post-column energy filter set to a slit width of 10 eV. The objective aperture was retracted and a 50 μm C2 aperture was inserted. Data were collected at pixel size of 0.84 Å pixels^−1^, a total dose of 60 e^−^ fractionated over 40 frames and a defocus range of −0.75 to –1.25 μm using EPU. The data set was collected at a dose rate of 33.914 e^−^ pixels^−1^ s^−1^. We acquired 5 images per hole and collected a total of 10510 micrographs.

Data were pre-processed using Warp (v.1.09) (Tegunov and Cramer, 2019). CTF parameters were estimated with a spatial resolution of 6 × 4 and motion correction was performed with a spatial resolution of 6 × 4. We picked 470,103 particles in Warp using a custom BoxNet model and extracted them in RELION (version 3.1) (Scheres, 2012) in a Box size of 672 Å. For initial classification, particles were binned to 3.42 Å pixel^−1^.

3D classification with six classes was performed on the extracted particles using a reference volume of a TREX–EJC–RNA on 15U reconstruction from a dataset collected on a Glacios TEM microscope low-pass filtered to 60 Å and a spherical mask of 550 Å diameter. Class 5 was selected with 84,300 octamer particles. To increase data set size, we separately extracted four THO–UAP56 dimers from each octamer, yielding a total of 329,826 dimers after removal of duplicates and re-extraction in CryoSPARC (box size 436 Å, 1.24 Å pixel^−1^). After another round of heterogeneous refinement with three classes the 204,147 particles of the best class where further refined through (1) a local refinement and non-uniform refinement using a TREX complex mask, yielding the 5.89 Å TREX complex Map A and (2) a local refinement using a mask including THO monomer 1A and UAP56, yielding the 4.12 Å THO–UAP56–ALYREF-N-UBM complex Map B.

#### Model building for the THO–UAP56–ALYREF-N-UBM complex

The structure of the human THO–UAP56 complex (PDB ID 7ZNL) (Pacheco-Fiallos et al., 2023) was docked into the THO–UAP56–ALYREF-N-UBM complex MapB. For UAP56, both RecA lobes were individually rigid-body fitted into the new map in COOT (Emsley and Cowtan, 2004; Emsley et al., 2010). The ALYREF N-UBM was fitted into the density based on an AlphaFold2 Multimer prediction of a UAP56–ALYREF complex and manually adjusted in COOT and the resulting structure was refined in phenix (Adams et al., 2010; Afonine et al., 2018) using the phenix.real_space_refine routine with secondary structure and rotamer restraints.

#### UAP56–UCM-1–N-UBM–TREX-2^M^ complex reconstitution and sample preparation

A PCID2–UAP56–UCM-1–N-UBM fusion protein in complex with SEM1 was combined with a 1.2x molar excess of MBP-GANP(582-1004) in buffer N (25 mM HEPES pH 7.9, 5% glycerol, 1 mM MgCl_2_, 1 mM TCEP, 200 μM 15U RNA, 1 mM AMP-PNP) and incubated on ice for 1 h. The sample was then centrifuged (21,130 x g, 15 min, 4°C) and applied to a Superdex 200 increase 10/300 size exclusion column, pre-equilibrated in buffer N, to separate the PCID2-UAP56–UCM-1–N-UBM–MBP-GANP(582-1004) complex from isolated components. Peak fractions were analyzed by SDS-PAGE and Coomassie staining to confirm stochiometric complex formation of the three proteins. The peak fraction was then diluted with buffer N to 0.8 mg ml^-1^ and cryoEM grids prepared by applying 4 µl of the sample to glow-discharged Cu R1.2/1.3 300-mesh holey carbon grids (Quantifoil). Grids were prepared blotted at 8 °C and 90% humidity and plunged into liquid ethane using a Leica EM GP2.

#### Cryo-EM data acquisition of a UAP56-UCM-1-N-UBM–TREX-2^M^ complex

Two datasets were collected on a 300 kV Titan Krios G4 equipped with a cold field emission gun, a post-column Selectris energy filter (ThermoFisher) with a 5 eV slit width and a Falcon 4i direct electron detector (ThermoFisher). The objective aperture was retracted and a 50 μm C2 aperture was inserted. For dataset 1 we collected 6,839 micrographs using EPU in the .eer format, with 5 images per hole, a pixel size of 0.749 Å pixel^-1^, a total does of 50 e^-^ / Å^2^ and a defocus range of −1 to −2.5 μm. Dataset 2 consists of 9,374 micrographs collected at a tilt angle of 20° and otherwise identical settings.

We performed on-the-fly preprocessing (patch motion correction and CTF estimation) using the CryoSPARC (Punjani et al., 2017) live routine. For dataset 1, we initially picked 3.8 mio particles in CryoSPARC live, extracted them with a 225 Å box and binned to 1.755 Å pixel^-1^ and performed 2 D classification. We then generated ab inito models for TREX-2^M^ and UAP56– TREX-2^M^, which were further refined through heterogeneous refinements and non-uniform refinements. These models were used as the initial reference maps for three rounds of heterogeneous refinement of 1.67 million particles picked in WARP and extracted with a 225 Å box and binned to 1.755 Å pixel^-1^, yielding 199,358 UAP56–TREX-2^M^ particles in the best class. These were re-extracted with a 225 Å box and binned to 0.877 Å pixel^-1^. Further heterogeneous refinement and 3D variability analysis yielded 19,188 UAP56–TREX-2^M^ particles in the best two classes.

For dataset 2 we picked 2.4 mio particles in WARP and extracted them with a 225 Å box and binned to 1.755 Å pixel^-1^. After 2D classification we obtained 660,903 TREX-2^M^ and UAP56–TREX-2^M^ particles. Two rounds of heterogeneous refinement yielded 316,490 TREX-2^M^ and 120,526 UAP56–TREX-2^M^ particles.

The 316,490 TREX-2^M^ particles were re-extracted with a 225 Å box and binned to 0.877 Å pixel^-1^, and subjected to a non-uniform refinement followed by 3D variability analysis. 57,499 particles from the best two clusters were refined through a local CTF refinement and a final local refinement with TREX-2^M^ mask yielded the 3.5 Å TREX-2^M^ complex Map C.

The UAP56–TREX-2^M^ particles were re-extracted with a 225 Å box and binned to 0.877 Å pixel^-1^ and subjected to another round of heterogeneous refinement, a non-uniform refinement, and a 3D variability analysis, resulting in 18,304 UAP56–TREX-2^M^ particles, which were combined with the UAP56–TREX-2^M^ particles from dataset 1. The combined 37,692 particles were subjected to a local CTF refinement and a final local refinement, leading to the 3.5 Å UAP56– TREX-2^M^ complex Map D. A further 3D classification with 20 classes and a GANP–UAP56-RecA2 mask, followed by local refinement of a class with 7,741 particles with bound UAP56-RecA2 lobe yielded the 4.22 Å UAP56–TREX-2^M^ complex Map E.

#### Model building for the TREX-2^M^ complex and the UAP56-UCM-1-N-UBM–TREX-2^M^ complex

An Alphafold2 Multimer prediction of TREX-2^M^ or UAP56–TREX-2^M^ was used as an initial model and docked into Map C and Map D densities, respectively. Model building was then manually adjusted in COOT and refined in phenix (Adams et al., 2010; Afonine et al., 2018) using the phenix.real_space_refine routine with secondary structure and rotamer restraints. The model of the UAP56 RecA2 lobe bound by the SARNP UCM-1 and the ALYREF N-UBM was obtained from an AlphaFold2 Multimer prediction and manually fitted into the UAP56 RecA2 density in Map E.

### ATPase assay

Steady-state UAP56 ATPase activity was measured using a NADH-coupled ATPase assay (Montpetit et al., 2012; Pühringer et al., 2020), with final concentrations of 5 U/mL rabbit muscle pyruvate kinase, Type III (Sigma-Aldrich), 5 U/mL rabbit muscle L-lactic dehydrogenase, Type XI (Sigma-Aldrich), 500 µM phosphoenolpyruvate and 50 µM NADH. Reactions were prepared in a final volume of 10 μl in a 1536 well plate and in buffer O (25 mM HEPES pH 7.9, 40 mM KCl, 0.5 mM MgCl_2_, 5% (w/v) glycerol, 0.5 mM ATP) with 0.5 / 2 µM UAP56 (when measured with TREX-2^M^ or in isolation), 2 µM TREX-2^M^ (wt or R678A mutant), 200 µM 15 U RNA. The NADH emission signal decay was monitored over time at 37°C in a PHERAstar FS (BMG LABTECH), with a 0.03–100 µM NADH dilution series as calibration standard. UAP56 ATPase rates were determined by linear regression of the NADH decay, corrected for ATP decay, as hydrolyzed molecules of ATP s^−1^ per enzyme. Input samples of the individual reactions were separated on 4 – 12% gradient SDS-PAGE gels and visualized by Coomassie staining.

### Generation of an endogenously tagged GFP–3C-UAP56 cell line

Human K562 cells (DSMZ) were edited to express an EGFP-3C-DDX39B fusion protein using a modification of a previously described CRISPR/Cas9 knock-in protocol (Koch et al., 2018).

Briefly, the gRNA was designed using the Benchling.com CRISPR gRNA design tool (Benchling; aaacTAACTGGGCCGGCAGGGGAAC) and cloned into the plasmid pLCG (hU6-sgRNA-EFSSpCas9-P2A-mCherry) (Muhar et al., 2018), a gift from J. Zuber, IMP, Vienna. The 500bp sequences flanking the DDX39B start codon were obtained by PCR on genomic K562 cDNA (using 5’ homology genomic primers: atcctcaagtaagggggtaccaggactctacttgtcatctccattttcc, gagatgttgaaggtcttcataactgggccagcagggga; and 3’ homology genomic primers: agggcccgggtggaggttccgctggagcagagaacgatgtggacaatg, atccccccttttcttttaaagaattctgatctagccttaagtataaaccc) and subcloned into pLPG vector (Muhar et al., 2018), a gift from J. Zuber, digested with MluI using Gibson Assembly (NEB), yielding the final vector pLPG-GFP-AID (5’ Blast_R_-P2A-eGFP-AID-3C).

K562 cells were grown in RPMI medium supplemented with 10% FBS (Sigma), 2% L-Glutamine (Gibco), 1% Sodium Pyruvate (Sigma) and 1% Penicillin Streptomycin (Sigma) and transfected with the HDR donor and the Cas9 plasmids using Neon electroporation device (Invitrogen) according to user guide manual (for suspension cells). 6 days post-transfection, after several passages, cells were subjected to fluorescence assisted cell sorting (FACS) using a BD FACSAria III (BD Biosciences). Cells expressing the EGFP-tag were sorted into 96 well plates. After approximately two weeks, wells with homogeneous fluorescence were genotyped (primers: TGCTAATTACACAAGGCTT, ACCTGCCACAGACCACTTCT), homozygous clones were further analysed by western blotting for homozygous knock-in of the tag using anti-UAP56 (ab181059; Abcam) and anti-GFP antibodies (A11122; Invitrogen).

### Export reporter

#### Generation of the K562 reporter cell line

The full reporter sequence, consisting of the mCherry coding sequence (CDS) with a single intron containing ten BoxB sites, an IRES and the GFP-Puromycin resistance ORF (mCherry^1/2^-5’SS-10xboxB-IRES-GFP-Puro^R^-3’SS-mCherry^2/2^), was synthesized (Genewiz) and cloned into a lentiviral vector backbone (Certo et al., 2011) (pRRL SFFV d20GFP.T2A.mTagBFP Donor was a gift from Andrew Scharenberg (Addgene plasmid # 31485 ; http://n2t.net/addgene:31485 ; RRID:Addgene_31485), yielding the plasmid containing pRRL-SFFV-reporter plasmid. Viral particles were generated by polyethylenimine transfection (Polysciences) of the pRRL-SFFV-reporter plasmid, together with the helper plasmids pCMVR8.74 (a gift from Didier Trono (Addgene plasmid # 22036 ; http://n2t.net/addgene:22036 ; RRID:Addgene_22036)) and pCMV-VSV-G (Stewart et al., 2003) (a gift from Bob Weinberg (Addgene plasmid # 8454 ; http://n2t.net/addgene:8454 ; RRID:Addgene_8454)) into LentiX-cells (Takara) according to standard procedures. K562 (DMSZ) cells were infected at limiting dilutions and mCherry positive single cells were isolated using a FACSAria III cell sorter (BD Biosciences). Viral integration of the entire reporter sequence was assessed by genotyping PCR. LentiX and K562 cells were maintained at 37 °C and 5% CO_2_ and tested negative for mycoplasma.

#### Plasmid transfection into K562 reporter cell line for lambdaN-mediated tethering

The coding sequence of a protein of interest was cloned into an acceptor plasmid containing the λN-BC2-FLAG tag, a P2A site and the BFP CDS (plasmid nLV-Ef1a, a gift from Stefan Ameres) using Gibson assembly (Gibson et al., 2009). For each protein of interest, a control plasmid lacking the λN-tag was created (Supplementary Table 5). Plasmids were transfected into the K562 reporter cell line using the Neon™ Transfection System 10 μL Kit (Invitrogen^TM^, cat.-Nr. MPK1025) according to the manual with 3 µg plasmid per 2×10^6^ cells (pulse voltage (V) =1,450, pulse width (ms) =10 and pulse number =3) in three replicates on different days. 48 h after transfection cells were analyzed using an iQue Screener Plus (Sartoriuos). Flow cytometry data was filtered for good events using FlowAI (Monaco et al., 2016), transfected K562 cells were selected by gating for BFP positive cells, and their GFP intensity extracted and plotted using GraphPad Prism (version 8).

To control for the expression and nuclear localization of λN-UAP56 and λN-UAP56 λNTD aliquots of one million cells were fractionated as previously described (Suzuki et al., 2010; Pacheco-Fiallos et al., 2023) and analyzed by westernblot using anti-UAP56 (ab181059, Abcam; 1:1000) and anti-Histone H3-HRP (5192S, Cell Signalling Technologies; 1:1000) antibodies.

### AlphaFold2 Multimer screening

Protein interaction prediction screening was performed using a custom pipeline (HT-Colabfold) based on Colabfold, which utilizes AlphaFold2 Multimer (Jumper et al., 2021; Evans et al., 2022; Mirdita et al., 2022). This pipeline was used to predict interactions between UAP56 and 696 proteins which were designated putative UAP56 interactors based on their at least two-fold enrichment over a wildtype control in UAP56-GFP immunoprecipitates.

HT-Colabfold manages the pairing, scheduling, and data collection for large-scale structure prediction and interaction screens. The pipeline executes pairwise predictions utilizing MMseqs 53 (git@92deb92) for local Multiple Sequence Alignment (MSA) generation (CPU-node) and Colabfold (git@7227d4c) for structure prediction (GPU-nodes). Each prediction involved the generation of 5 models, omitting structure relaxation. Predictions with an average iPTM score of > 0.5 were considered putative hits and diagnostic plots (PAE plot, pLDDT plot and sequence coverage) as well as the generated structures were manually inspected.

### Code availability

HT-Colabfold is free open-source software (MIT) and available at https://gitlab.com/BrenneckeLab/ht-colabfold.

## SUPPLEMENTARY FIGURES

**Supplementary Figure 1.**
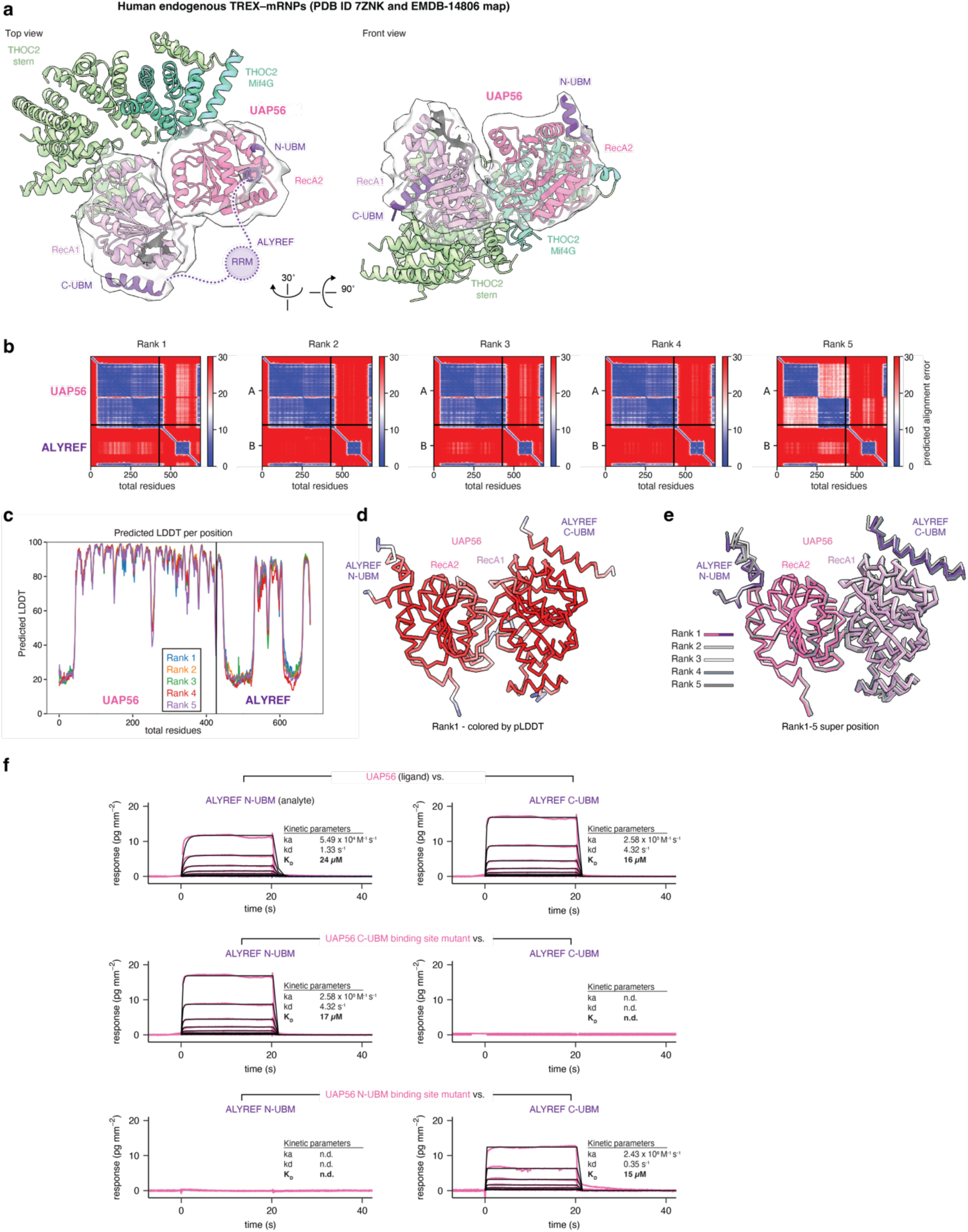
ALYREF N- and C-UBMs bind distinct UAP56 surfaces. **a,** Re-analysis of a human endogenous TREX**–**mRNP structure reveals an unidentified binding site for a N-UBM on UAP56. Front (left) and top views (middle) of UAP56 bound to THO and mRNPs with the THOC2 MIF4G and Stern domains (green), UAP56 (pink) and the ALYREF C-UBM (purple) (PDB ID 7ZNK), with cryo-EM density around UAP56 and ALYREF shown (EMDB-14806). An additional density at low resolution on UAP56’s RecA2 lobe fits with the AlphFold2 predicted binding site of ALYREF’s N-UBM (purple). **b-e,** Diagnostic plots for the AlphaFold2 Multimer prediction of ALYREF and UAP56. The N- and C-UBM are predicted with high confidence to bind distinct binding sites: the C-UBM is predicted in the previously experimentally determined binding site on the RecA1 lobe, the N-UBM on a novel binding site on RecA2. Shown are **b,** the PAE plot, **c,** the pLDDT plot, **d,** the structure of the top ranked model in Cα trace colored by pLDDT (shown are only the ordered and interacting elements: UAP56 RecA1 and RecA2, residues 40-428; ALYREF N-UBM and C-UBM, residues 1-24 and 236-257), and **e,** a superposition of the structures of all five models, rank 1-5, as in **d,** but colored for rank 1 with UAP56 in pink and ALYREF in purple, and in shades of grey for rank 2-5. **f,** Biochemical evidence that the N-UBM binds a binding site on UAP56 distinct from the C- UBM. GCI derived direct binding kinetics for UBM peptides as the analyte and UAP56 as the ligand (see Fig. 1d). Sensograms (pink), fit (black) with table summaries shown alongside. UAP56 binds N- and C-UBM peptides with mid-micromolar affinity (top row). Mutating key residues in the UAP56 C-UBM binding site (R208S, Q212A, R216S, K241S; RQRK) prevents C-UBM binding but leaves N-UBM binding unaffected (middle row). Mutating key residues in the newly identified N-UBM binding site (R276E, A302R, E309S) abolishes N-UBM binding without affecting C-UBM binding (bottom row).

**Supplementary Figure 2.**
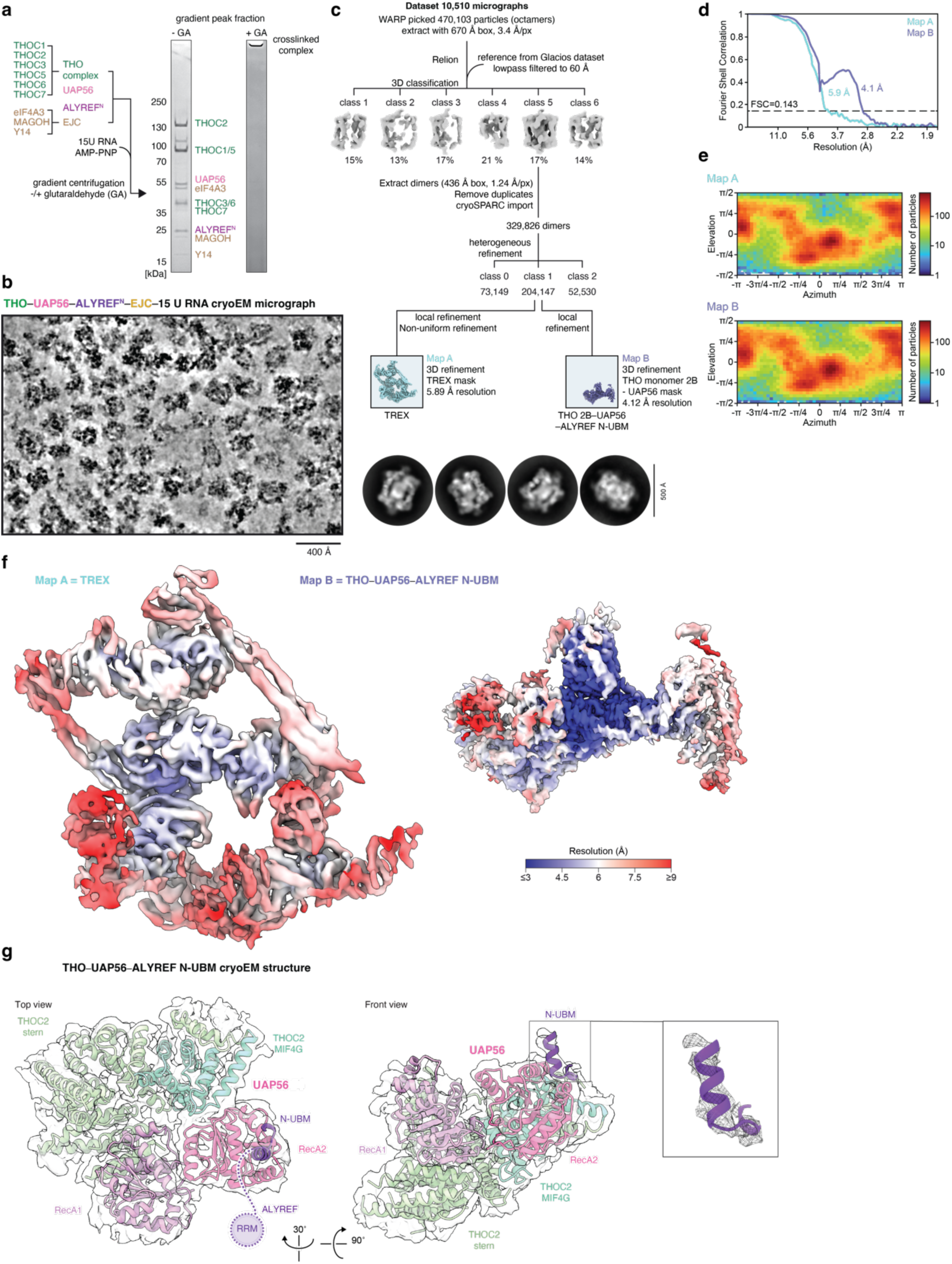
TREX–EJC–RNA complex reconstitution and cryo-EM analysis. **a,** Complex reconstitution scheme for TREX (=THO–UAP56–ALYREF^N^)–EJC–RNA from recombinant proteins. Complex components were mixed and separated on a 15-40 % sucrose gradient with or without 0.05 % glutaraldehyde, fractionated and analyzed by Coomassie-stained SDS-PAGE. Shown are peak lanes of the crosslinked complex used for cryo-EM sample preparation (right) the corresponding fraction from the non-crosslinked gradient (left). **b,** Denoised cryo-EM micrograph of TREX–EJC–RNA. Scale bar, 400 Å. **c,** Three-dimensional image classification tree, with representative 2D classes shown below. The dataset contains 10,510 micrographs, of which 470,103 particles (THO octamer) were picked with WARP (Tegunov and Cramer, 2019) and processed in Relion (Scheres, 2012) and cryoSPARC (Punjani et al., 2017). The final particle stack contained 204,147 particles and was refined to 5.89 Å for the entire TREX complex (Map A) and to 4.12 Å for a THO monomer (Map B). **d,** Gold-standard Fourier shell correlation (FSC = 0.143) of the TREX complex (Map A) and THO-monomer–UAP56–N-UBM (Map B) cryo-EM maps. **e,** Orientation distribution plots, as visualized in cryoSPARC, for all particles contributing to the TREX complex (Map A) and THO-monomer–UAP56–N-UBM (Map B) cryo-EM maps. **f,** TREX complex (Map A) and THO-monomer–UAP56–N-UBM (Map B) cryo-EM maps colored by local resolution. **g,** Structure of the THO-monomer–UAP56–N-UBM complex (left, top view; middle and right, front view), with the THOC2 MIF4G and Stern domains (green), UAP56 (pink) and the N-UBM (purple) shown together with the Map B density (inset on the right). The structure reveals density for the ALYREF N-UBM at the newly identified binding site in the UAP56 RecA2 lobe.

**Supplementary Figure 3.**
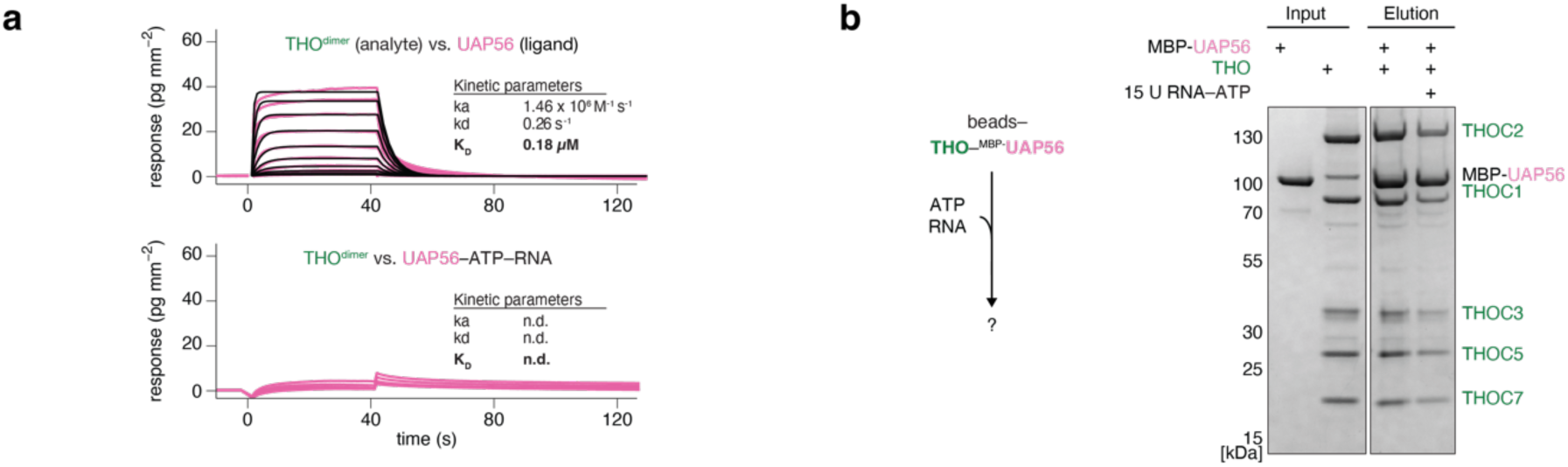
ATP and RNA alone do not dissociate the THO–UAP56 complex. **a,** GCI derived direct binding kinetics for a THO dimer as the analyte and UAP56 as the ligand (compare to Fig. 1d). Sensograms (pink), fit (black) with table summaries shown alongside. **b,** ATP and RNA lead to the partial dissociation of THO from UAP56 *in vitro*. Experimental setup (left) and Coomassie stained SDS-PAGE gel of a representative experiment (right).

**Supplementary Figure 4.**
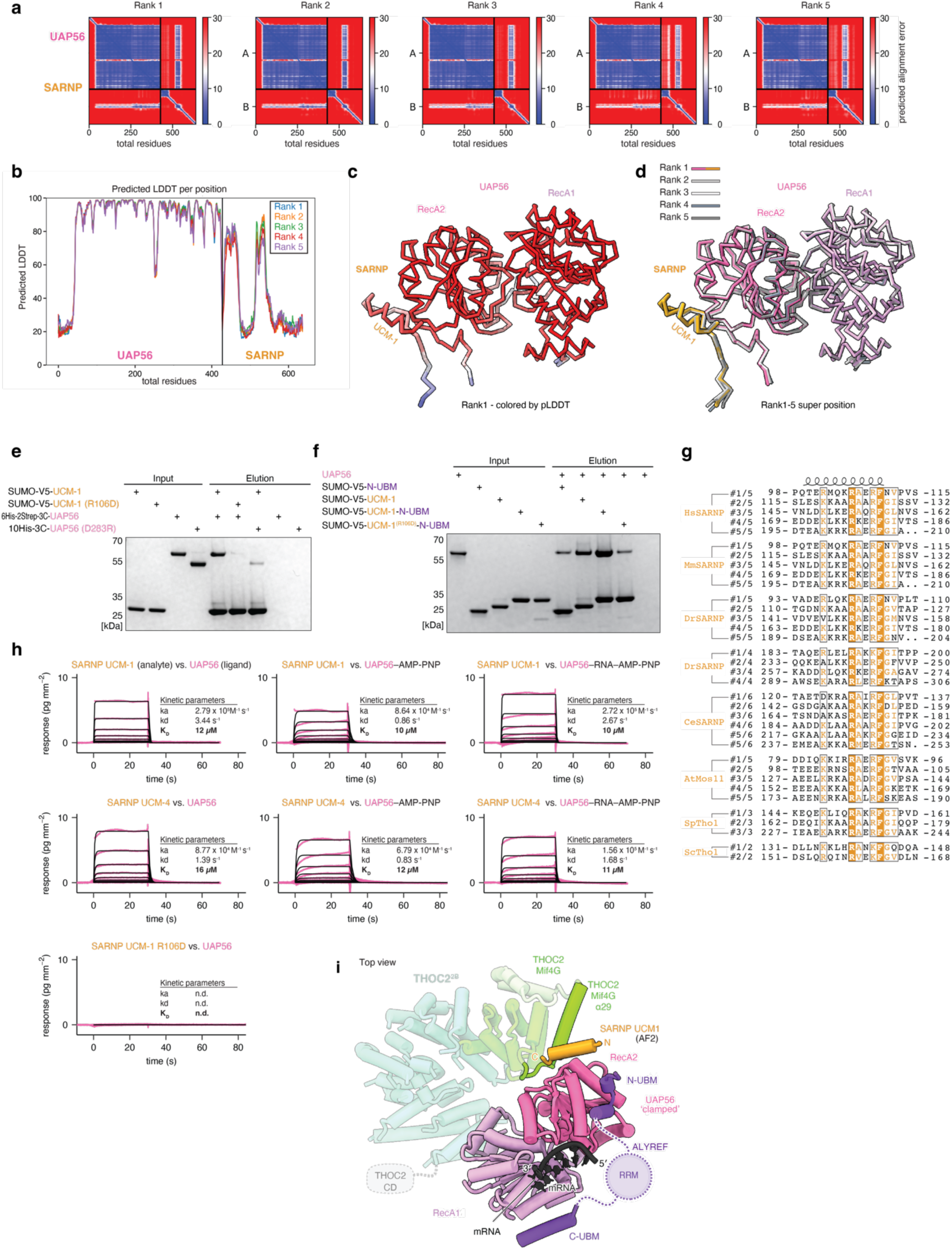
SARNP UCMs bind UAP56. **a-d,** Diagnostic plots for the AlphaFold2 Multimer prediction of SARNP–UAP56. A UCM is predicted with high confidence to bind a binding site on UAP56’s RecA2 lobe. Shown are **a,** the PAE plot, **b,** the pLDDT plot, **c,** the structure of the top ranked model in Cα trace colored by pLDDT (shown are only the ordered and interacting elements: UAP56 RecA1 and RecA2, residues 40-428; SARNP UCM-1 region, residues 84-110), and **d,** a superposition of the structures of all five models, rank 1-5, as in **c,** but colored for rank 1 with UAP56 in pink and SARNP in yellow, and in shades of grey for rank 2-5. **e,** UAP56 interacts with a SARNP UCM. Recombinant SUMO-V5 tagged SARNP UCM-1 (residues 82-115) is immobilized on magnetic V5 beads and incubated with recombinant full length UAP56. After washing out unbound UAP56, bead-bound complexes are eluted at low pH and analyzed by SDS-PAGE followed by Coomassie staining, revealing that UAP56 interacts with the UCM-1 (lane 5). Mutating key residues in the predicted interface (SARNP UCM-1 R106D or UAP56 D283R) prevents the interaction almost completely (lanes 6 and 7). UAP56 shows little background binding (lanes 8 and 9). **f,** SARNP UCM-1 and the ALYREF N-UBM can bind UAP56 simultaneously. SUMO-V5-3C–N-UBM, SUMO-V5-3C–UCM, or a SUMO-V5-3C–UCM–N-UBM fusion (with or without the SARNP R106D) mutation are immobilized on magnetic V5 beads, incubated with UAP56 and washed, eluted and analyzed as in **e,**. UAP56 binds the isolated UCM-1 with higher affinity than the N-UBM (lanes 6 and 7). The binding affinity to the UCM–N-UBM fusion is higher than for the individual peptides (lane 8 vs. lanes 6 and 7), suggesting that both peptides can bind simultaneously. Mutating R106 in the UCM–N-UBM fusion reduces UAP56 binding to the level of the N-UBM alone (lane 9), and UAP56 shows little background binding (lane 10). **g,** Multiple sequence alignment of the SARNP UCMs from *H. sapiens* (Uniprot ID P82979), *M. musculus* (Mm, Q9D1J3), *D. rerio* (Dr, Q504C3), *D. melanogaster* (DmCG8149, Q9VHC8), *C. elegans* (Ce, Q9N3G0), *A. thaliana* (AtMos11, Q9LZ08), *S. pombe* (Sp, O74871) and *S. cerevisiae* (ScTho1, P40040) (bottom), colored by conservation (orange underground = invariant residue). **h,** GCI derived direct binding kinetics for a SARNP UCM peptides as the analyte and UAP56 as the ligand. UAP56 binds the SARNP UCM-1 (top row) and UCM-4 (middle row) with similar affinity (∼ 10 to 20 μM), and binding is affected neither by the addition of the non-hydrolysable ATP analogue AMP-PNP nor by AMP-PNP and 15 poly-Uridine RNA. The mutation R106D in UCM-1 prevents UAP56 binding (bottom). Sensograms (pink), fit (black) with table summaries shown alongside. **i,** Structural modeling of a UAP56–RNA–SARNP-UCM-1–ALYREF-N-UBM complex (see Fig. 2d) superimposed on THO–UAP56 reveals a clash between the SARNP UCM and the MIF4G domain in THOC2. UAP56, pink; RNA, black; SARNP, yellow; ALYREF UBMs, purple; THOC2 MIF4G and Stern in green and, except for the clashing α-helix 29, transparent.

**Supplementary Figure 5.**
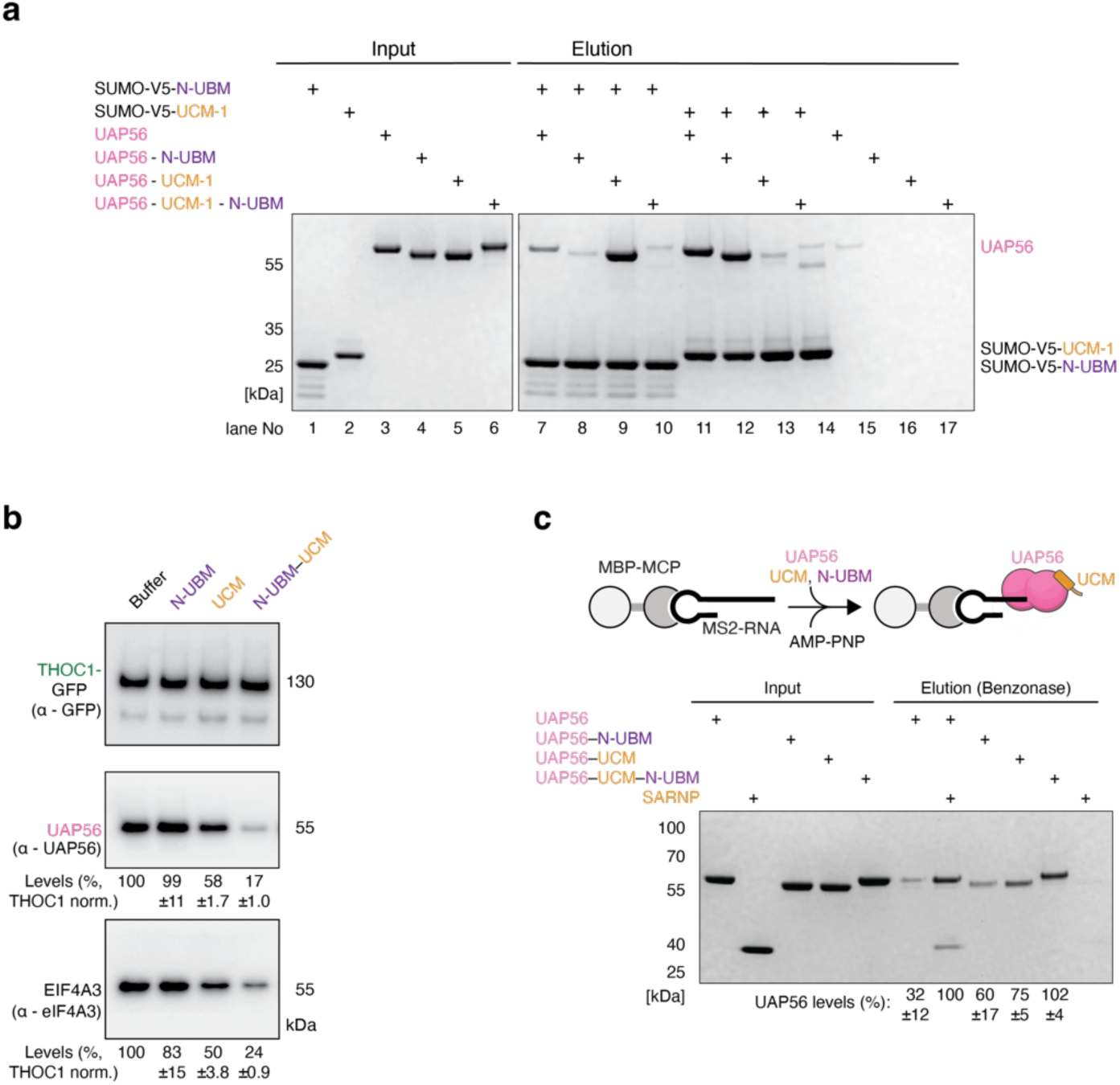
ALYREF N-UBM and SARNP UCM cooperatively bind UAP56 and SARNP assists TREX disassembly and stabilizes mRNA-clamped UAP56. **a,** SUMO-V5-3C tagged N-UBM or UCM-1 are immobilized on magnetic V5 beads and incubated with UAP56 or UAP56–N-UBM, UAP56–UCM or UAP56–UCM–N-UBM fusion proteins. Beads were washed, eluted, and analyzed as in Supplementary Fig. 4e. UAP56 binds both N-UBM and UCM (lanes 7 and 11). The UAP56–N-UBM fusion binds the UCM like wildtype UAP56, but not the N-UBM (lanes 12 and 8). UAP56–UCM does not bind to immobilized UCM and binds the N-UBM with apparent higher affinity than UAP56 alone (lanes 13 and 9), suggesting that N-UBM and UCM bind cooperatively. The UAP56–UCM– N-UBM fusion binds neither immobilized N-UBM nor UCM (lanes 10 and 14), suggesting that both peptides are bound to their cognate binding site in the fusion protein. None of the UAP56 proteins exhibits relevant background binding (lanes 15-18). **b,** Western blot analysis for the endogenous TREX disassembly experiment depicted in Fig. 2f. **c,** UAP56 is stabilized on RNA by SARNP. A 450 nt long MS2-loop containing AdML RNA is immobilized on amylose beads through an MBP-MCP fusion protein. The RNA is incubated in the presence of the non-hydrolysable ATP analogue AMP-PNP with UAP56, UAP56 and full length SARNP or with UAP56–N-UBM, UAP56–UCM or UAP56–UCM– N-UBM fusion proteins. Unbound proteins are washed out, and RNA-bound proteins are eluted through benzonase digestion of the RNA and visualized on Coomassie stained SDS-PAGE gels, with the quantification of bound UAP56 shown alongside (UAP56 with SARNP is set to 100 % bound). UAP56 alone shows little RNA binding. The presence of a N-UBM or UCM increases the RNA bound fraction, and the presence of the UCM–N-UBM fusion or full length SARNP leads to maximum RNA binding.

**Supplementary Figure 6.**
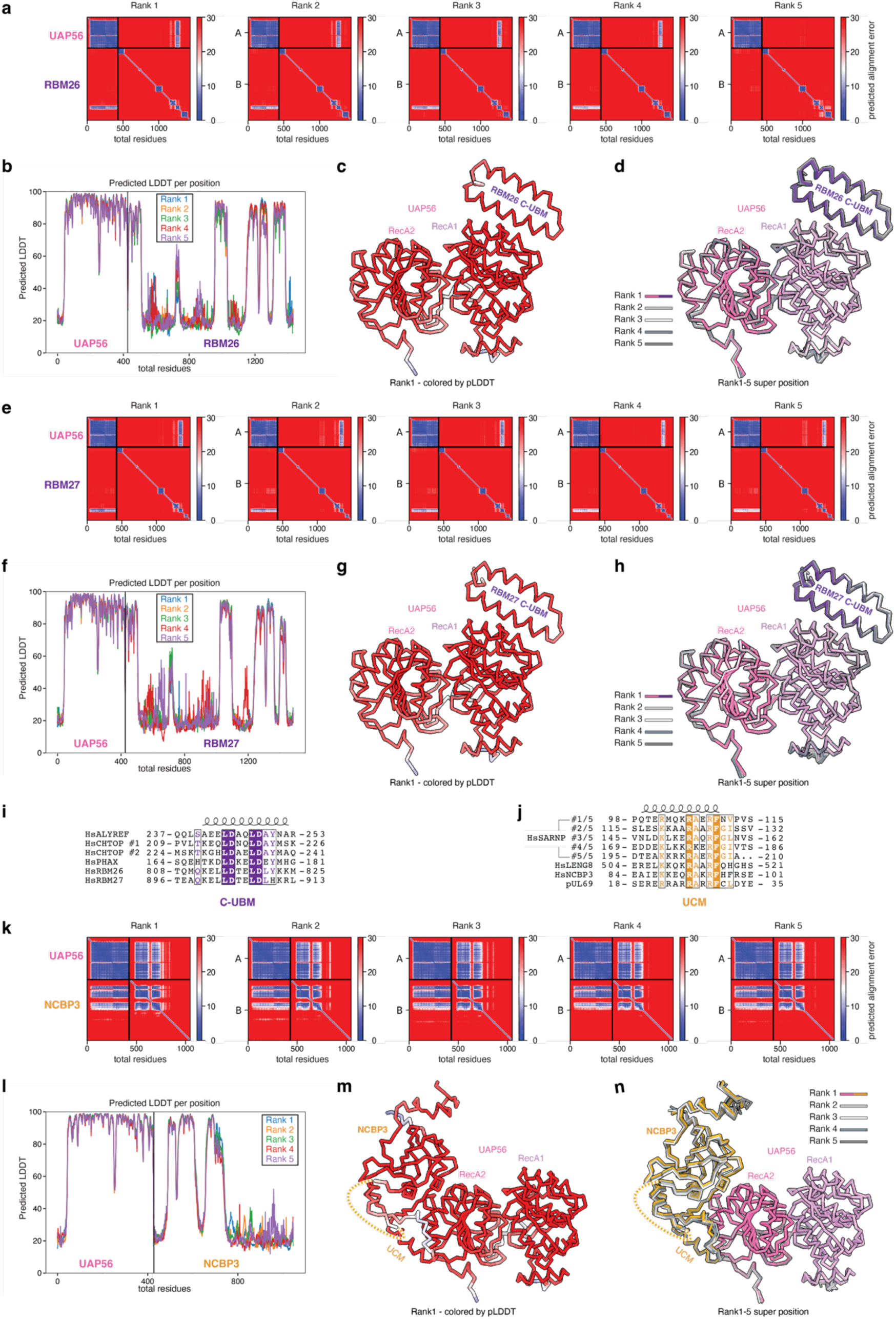
AlphaFold2 Multimer identifies putative UAP56 interactors. **a-h,** Diagnostic plots for the AlphaFold2 Multimer prediction of the paralogs RBM26 **a-d,** or RBM27 **e-h,** and UAP56. A putative C-UBM is predicted for RBM26 and RBM27 with high confidence to bind the characterized C-UBM binding site on the RecA1 lobe of UAP56. Shown are **a,e,** the PAE plots, **b,f,** the pLDDT plots, **c,g,** the structures of the top ranked model in Cα trace colored by pLDDT (shown are only the ordered and interacting elements: UAP56 RecA1 and RecA2, residues 40-428; RBM26/27 C-UBM region residues 803-852 / 891-940), and **d,h,** a superposition of the structures of all five models, rank 1-5, as in **c,g,** but colored for rank 1 with UAP56 in pink and RBM26/27 in purple, and in shades of grey for rank 2-5. **i,** Multiple sequence alignment of known and novel human C-UBM containing proteins (HsALYREF, Uniprot ID Q86V81, HsCHTOP Q9Y3Y2, HsPHAX Q9H814, HsRBM26 Q5T8P6, HsRBM27 Q9P2N5), colored by conservation (purple underground = invariant residue). **j,** Multiple sequence alignment of putative UCM motifs in human SARNP, LENG8, NCBP3 and the Human cytomegalovirus protein pUL69 (HsSARNP, Uniprot ID P82979, HsLENG8 Q96PV6, HsNCBP3 Q53F19, pUL69 P16749), colored by conservation (orange underground = invariant residue). **k-n,** Diagnostic plots for the AlphaFold2 Multimer prediction of NCBP3 and UAP56. A UCM containing region is predicted with high confidence to bind to the RecA2 lobe of UAP56. Shown are **k,** the PAE plot, **l,** the pLDDT plot, **m,** the structure of the top ranked model in Cα trace colored by pLDDT (shown are only the ordered and interacting elements: UAP56 RecA1 and RecA2, residues 40-428; NCBP3, residues 59-184 and 231-295), and **n,** a superposition of the structures of all five models, rank 1-5, as in **m,** but colored for rank 1 with UAP56 in pink and NCBP3 in yellow, and in shades of grey for rank 2-5.

**Supplementary Figure 7.**
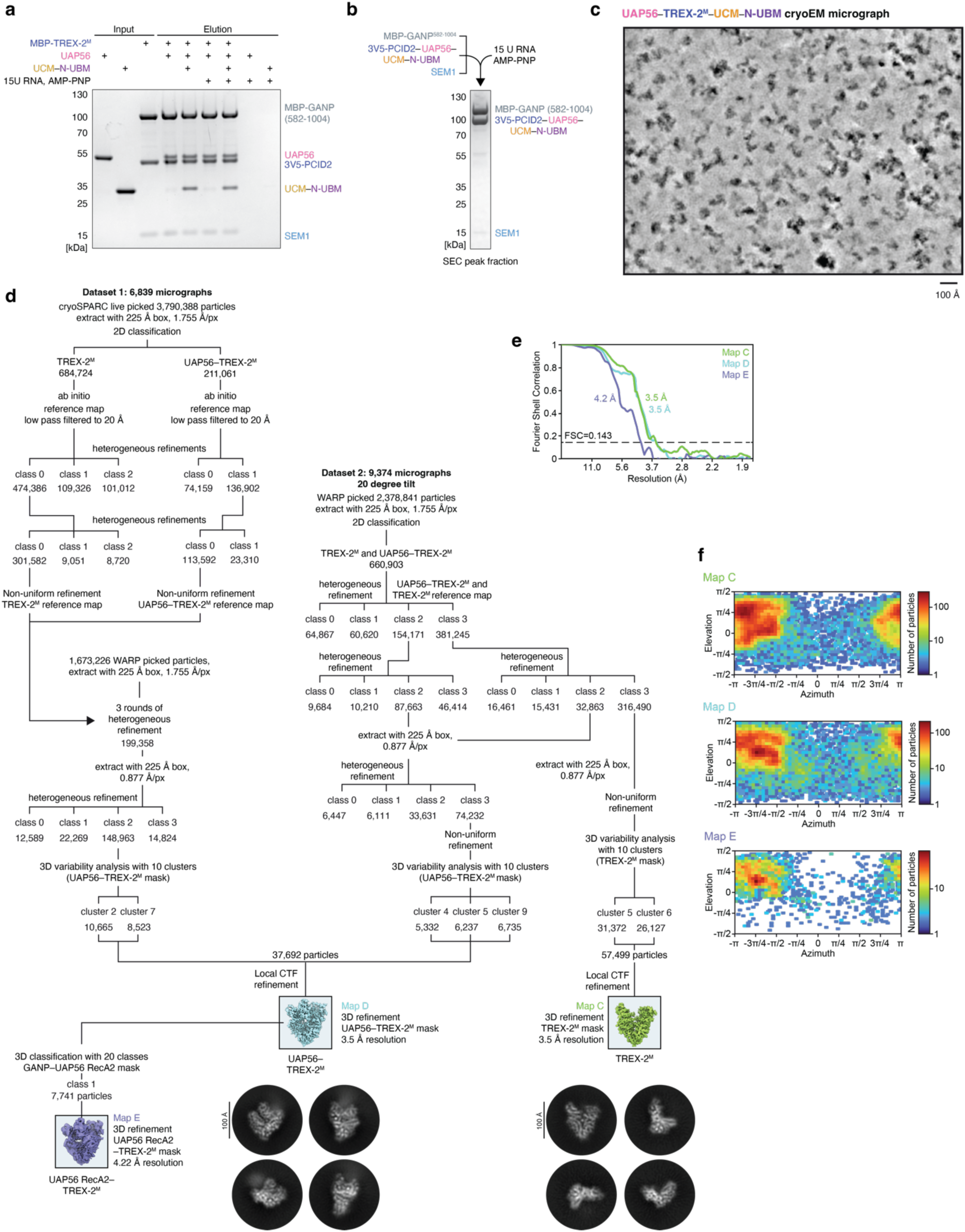
TREX-2^M^ and UAP56–TREX-2^M^ complex cryo-EM analysis. **a,***In vitro* pulldown assay probing the UAP56–TREX-2^M^ interaction. Recombinant TREX-2^M^ complex (MBP-GANP residues 582-1004, PCID2, SEM1) is immobilized on amylose beads through the MBP on the GANP subunit and incubated with UAP56 in the presence or absence of UCM–N-UBM peptide and / or 15 U RNA and AMP-PNP. After washes the bead bound proteins are eluted with a maltose containing buffer and visualized in Coomassie stained SDS-PAGE gels. UAP56 forms a near stochiometric complex with TREX-2^M^, and complex formation is compatible with binding of the UCM–N-UBM peptide and the presence of 15 U RNA and AMP-PNP. **b,** Complex reconstitution scheme for the UAP56–TREX-2^M^ complex for cryo-EM. MBP-GANP residues 582-1004 was incubated with a complex consisting of the PCID2–UAP56– UCM–N-UBM fusion protein and SEM1, 15 poly-Uridine RNA and the non-hydrolysable ATP analogue AMP-PNP and the formed complex separated by size exclusion chromatography, with a peak fraction shown on an Coomassie-stained SDS-PAGE gel. **c,** Denoised cryo-EM micrograph of the UAP56–TREX-2^M^ sample. Scale bar, 100 Å. **d,** Three-dimensional image classification tree, with representative 2D classes shown below. From two datasets, containing 6,839 and 9,374 (collected at 20-degree stage tilt) micrographs, 1,673,226 and 2,378,841 particles were picked using a custom trained BoxNet in WARP and processed in cryoSPARC. The final particle stacks contained 57,499 particles for TREX-2^M^ (Map C) and 37,692 particles for UAP56–TREX-2^M^ (Map D) and were each refined to 3.5 Å resolution. The Map D particle stack was further classified using a GANP– UAP56 RecA2 mask, yielding a particle stack of 7,741 particles with observable RecA2 density which was refined to 4.22 Å resolution (Map E). **e,** Gold-standard Fourier shell correlation (FSC = 0.143) of the TREX-2^M^ complex (Map C), UAP56 RecA1–TREX-2^M^ complex (Map D), and the UAP56 RecA1-RecA2–TREX-2^M^ complex (Map E) cryo-EM maps. **f,** Orientation distribution plots, as visualized in cryoSPARC, for all particles contributing to the TREX-2^M^ complex (Map C) the UAP56 RecA1–TREX-2^M^ complex (Map D), and the UAP56 RecA1-RecA2–TREX-2^M^ complex (Map E) and cryo-EM maps.

**Supplementary Figure 8.**
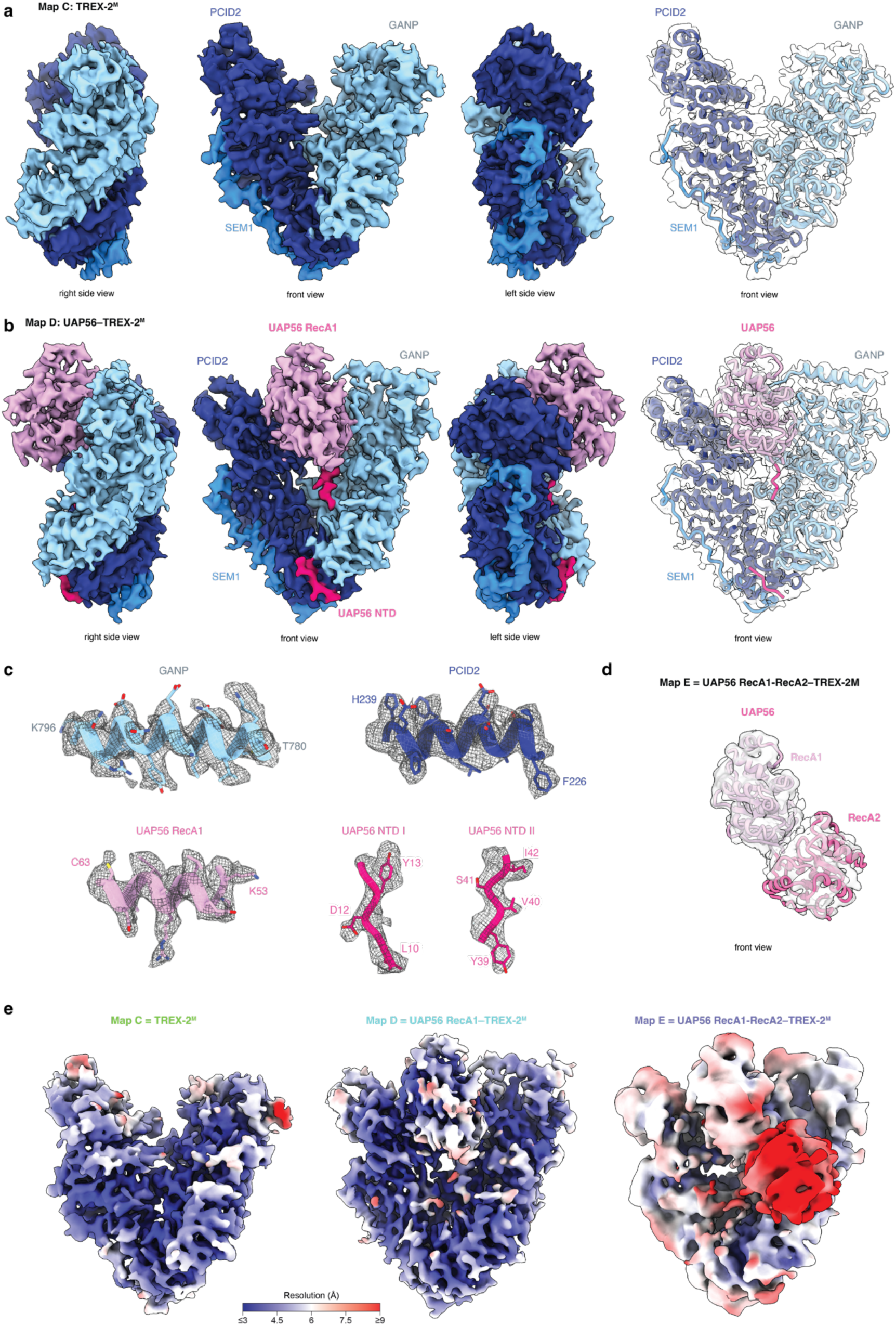
TREX-2^M^ and UAP56–TREX-2^M^ complex cryo-EM analysis. **a,**TREX-2^M^ complex (Map C) cryo-EM density, shown in left, front, and right side views and colored by subunit (GANP, light blue; PCID2, dark blue; SEM1, dodger blue). Shown alongside on the very right is the superposition of the TREX-2^M^ model, in cartoon representation and again colored by subunit, superimposed on the cryo-EM Map E. **b,** UAP56 RecA1–TREX-2^M^ complex (Map D) cryo-EM density, shown in left, front, and right side views and colored by subunit (UAP56 RecA1, pink; GANP, light blue; PCID2, dark blue; SEM1, blue). Shown alongside on the very right is the superposition of the UAP56 RecA1–TREX-2^M^ model, in cartoon representation and again colored by subunit, superimposed on the cryo-EM Map C. **c,** Representative segments of GANP, PCID2 and UAP56 NTD and RecA1 from Map D superimposed on the respective cryo-EM densities. **d,** Superposition of the UAP56 RecA1 RecA2 model, in cartoon representation and colored in pink, on the cryo-EM Map E. **e,** TREX-2^M^ complex (Map C), UAP56 RecA1–TREX-2^M^ complex (Map D), and UAP56 RecA1 RecA2–TREX-2^M^ complex (Map E) cryo-EM maps colored by local resolution.

**Supplementary Figure 9.**
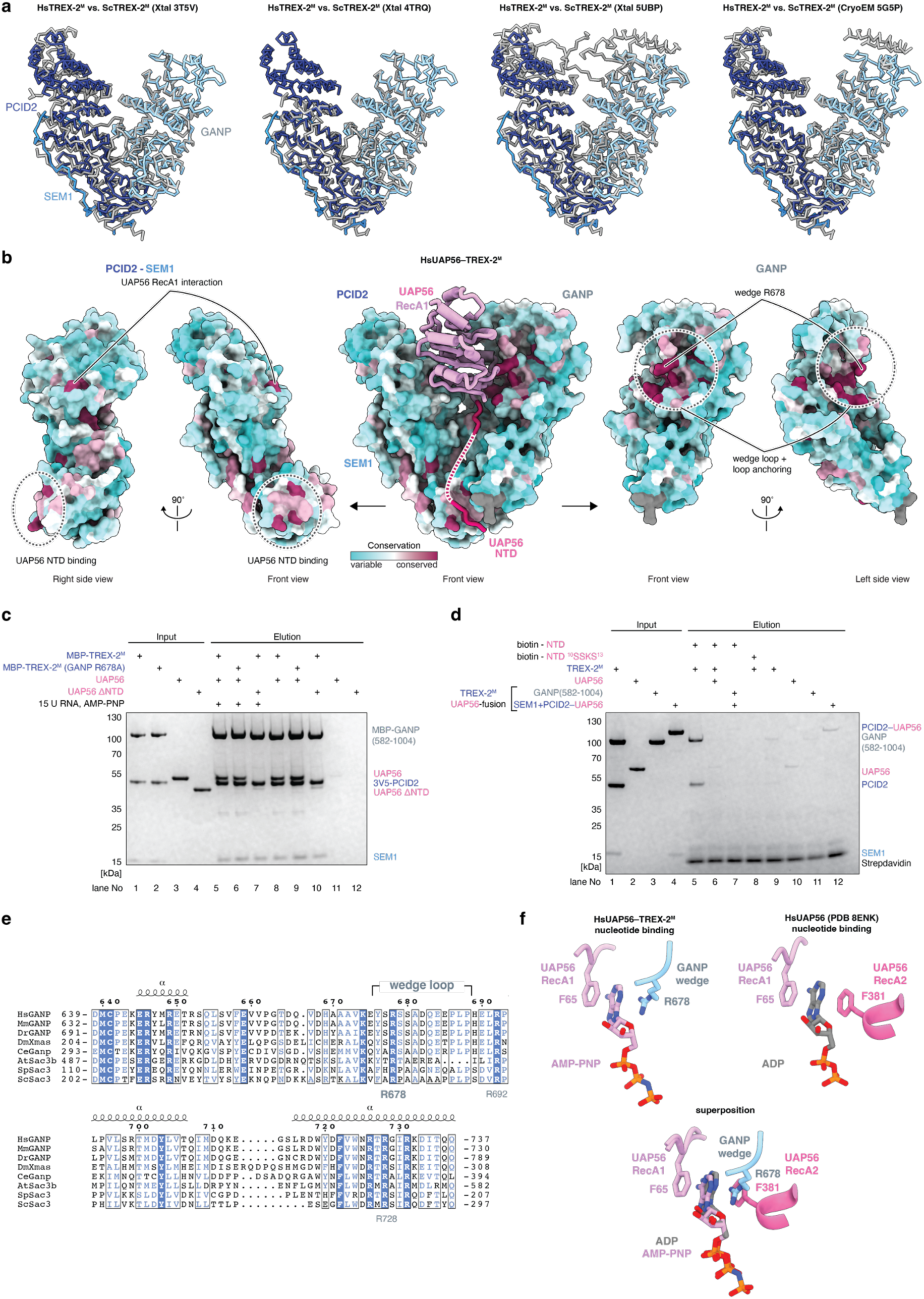
Structural analysis of TREX-2^M^ and UAP56–TREX-2^M^. **a,** The human TREX-2^M^ complex cryo-EM structure is highly similar to previous crystal structures and a cryo-EM structure of the yeast TREX-2^M^ complex. Shown are superpositions of the human TREX-2^M^ model determined in this study, in front view and as Cα trace colored by subunit (GANP, light blue; PCID2, dark blue; SEM1, dodger blue), on previous structures of the yeast TREX-2^M^ complex (in grey) determined through crystallography (left, PDB ID 3T5V; middle left, PDB ID 4TRQ; middle right, PDB ID 5UBP) or cryo-EM (right, PDB ID 5G5P) (Ellisdon et al., 2012; Schneider et al., 2015; Aibara et al., 2016; Gordon et al., 2017). **b,** Structure of the UAP56**–**TREX-2^M^ complex, with UAP56 in pink and cartoon representation and PCID2, SEM1 and GANP in surface representation and colored by sequence conservation (maroon, conserved; cyan, variable). A front view of the entire complex is shown in the center, flanked by front and side views of PCID2**–**SEM1 (left) or GANP (right). Surfaces patches involved in complex formation, such as the UAP56 RecA1 proximal patch in PCID2, the UAP56 NTD binding site, the GANP wedge or the PCID2**–** GANP interface, show a high degree of sequence conservation. **c,** *In vitro* pulldown assay as in Supplementary Fig. 7a, but using wildtype TREX-2^M^ or the wedge mutant TREX-2^M^ (containing GANP R678A) and wildtype UAP56 or UAP56 ΔNTD (residues 44-428). A UAP56–TREX-2^M^ complex is formed with the wedge loop mutant TREX-2^M^, while the deletion of the UAP56 NTD abolishes complex formation. **d,** *In vitro* pulldown assay probing TREX-2^M^ binding of the isolated UAP56 NTD. Biotinylated UAP56 NTD peptides (residues 1-21, wildtype or mutant E9K, L10S, L11A, D12K, Y13S) are immobilized on streptavidin beads and incubated with TREX-2^M^ in the presence or absence of UAP56, or with TREX-2^M^ with UAP56 fused to the C-terminus of PCID2. Protein complexes are eluted and visualized on Coomassie stained SDS-PAGE gels. Wildtype, but not mutant, NTD peptide forms a complex with TREX-2^M^, and complex formation is abolished in the presence of full length UAP56. TREX-2^M^ with UAP56 fused to the PCID2 C-terminus also does not form a complex with NTD peptide, suggesting that UAP56 is TREX-2^M^-bound in the fusion construct. **e,** Multiple sequence alignment of the GANP wedge loop region of GANP proteins from *H. sapiens* (HsGANP, Uniprot ID O60318), *M. musculus* (MmGANP, Q9WUU9), *D. rerio* (DrGANP, F1Q712), *D. melanogaster* (DmXmas, Q9U3V9), *C. elegans* (CeGanp, Q19643*), A. thaliana* (AtSac3b, F4JAU2), *S. pombe* (SpSac3, O74889) and *S. cerevisiae* (ScSac3, P46674), colored by conservation (blue underground = invariant residue), and with secondary structure elements depicted on top. Highlighted are the wedge loop with the key and invariant residue R678 (numbering according to HsGANP), as well as the invariant residues R692 and R728 which might stabilize the wedge loop and UAP56 RecA1 interactions and are implicated in gene gating (Schneider et al., 2015). **f,** The GANP wedge binding mimics the ATP-binding pocket in clamped UAP56. Shown are the UAP56-bound nucleotide and nucleotide base stacking residues in sticks representation, colored by heteroatom, and with UAP56 in pink, GANP in light blue and the nucleotide in pink for the UAP56**–**TREX-2^M^ structure (left), and in grey for the clamped human UAP56 structure (right, PDB ID 8ENK) (Xie et al., 2023). In the UAP56**–**TREX-2^M^ structure the adenine moiety of the non-hydrolyzable ATP analogue forms stacking interactions with F65 in the UAP56 RecA1 lobe and the GANP wedge residue R678 (left). In clamped UAP56, the adenine base of bound ADP stacks with F65 of UAP56’s RecA1 lobe and F381 of the RecA2 lobe (right). Superposition of both structures reveals that the GANP wedge residue R678 substitutes the RecA2 lobe residue F381 (bottom).

**Supplementary Figure 10.**
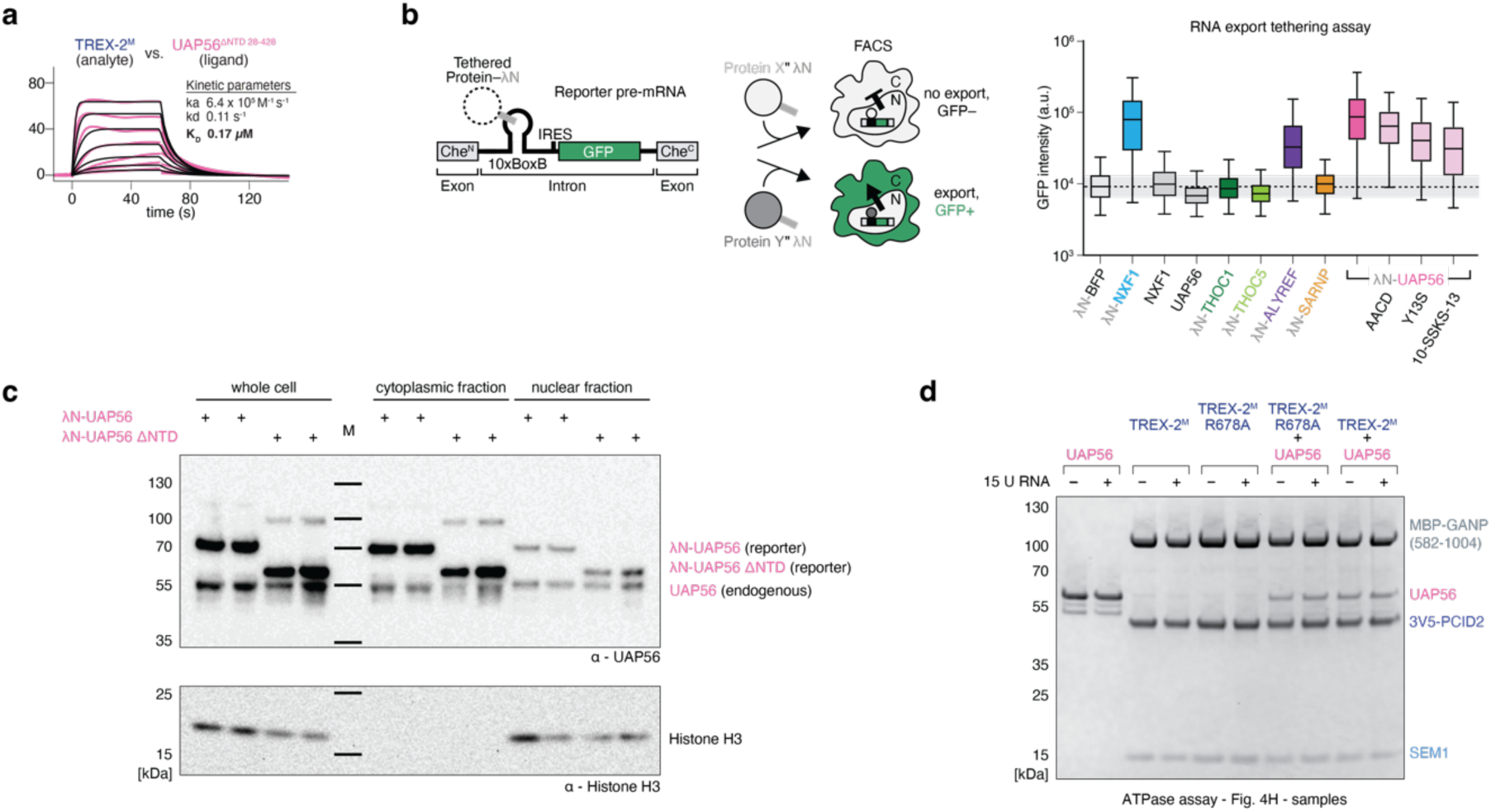
Controls for RNA export reporter and UAP56 ATPase assays. **a,** Sensogram for TREX-2^M^ (immobilized) probed with UAP56ΔNTD (residues 28-428). Sensogram (pink line), the fitted model (black), and a binding kinetics summary table are shown. Related to Fig. 4d. **b,** RNA export tethering assay, related to Fig. 4e with λN-BFP, λN-NXF1 and λN-UAP56 from Fig. 4e shown for comparison. Mutations interfering with UAP56’s ATPase activity (D196A, E197A, ‘AACD’) do not substantially alter the export promoting effect, while mutating key residues in the UAP56 NTD (Y13S or L10S, L11S, D12K, Y13S, ’10-SSKS-13’) leads to a reduced effect. As additional controls, we also tethered the THO complex via THOC1 or THOC5, or of SARNP, which showed no effect, as well as ALYREF, which modestly stimulated reporter pre-mRNA export. **c,** λN-UAP56 and λN-UAP56 ΔNTD are expressed and imported to the nucleus at similar levels. Two replicates of λN-UAP56 or λN-UAP56 ΔNTD expressing K562 cells are fractionated into nucleus and cytoplasm, proteins separated by SDS-PAGE and analysed by western blotting, probing for UAP56 (top) and Histone H3 (bottom, fractionation control) on the same membrane; the full membrane is shown. Both constructs are expressed to similar levels (as judged from the whole cell extract, compared to endogenous UAP56 to showcase equal loading). Equal amounts of each construct are important into the nucleus, and nuclear levels of the λN-tagged proteins are comparable to levels of endogenous nuclear UAP56. **d,** Samples used in the ATPase assay in Fig. 4f. An aliquot of each reaction was separated in SDS-PAGE and visualized by Coomassie staining.

**Supplementary Table 1.**
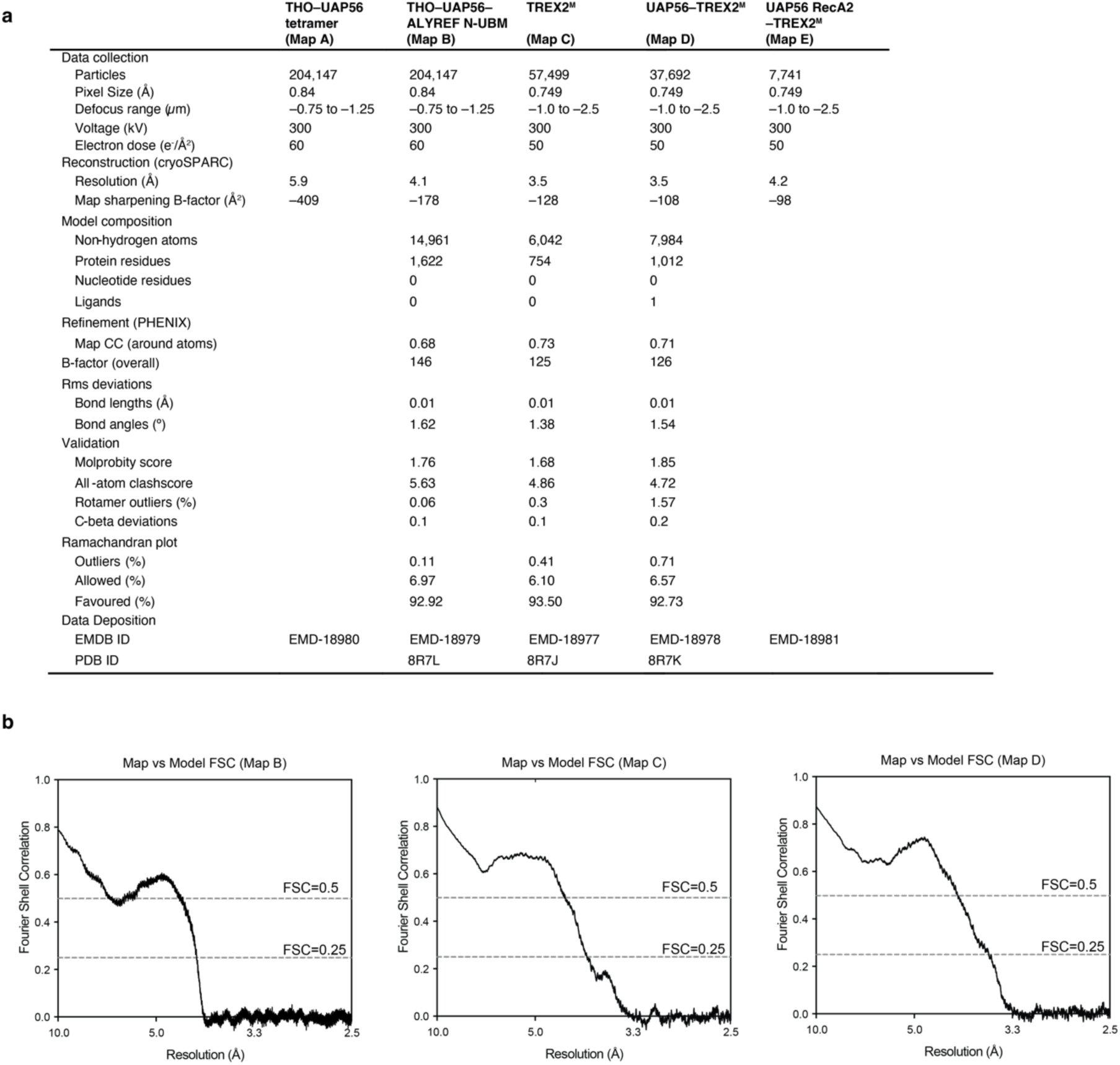
Cryo-EM data collection and refinement statistics. **a,** Cryo-EM data collection and refinement statistics for the THO–UAP56–ALYREF-N-UBM complex as well as TREX-2^M^ and UAP56–TREX-2^M^ **b,** Fourier shell correlations between the cryo-EM densities of the THO–UAP56–ALYREF-N-UBM complex (Map B, left), TREX-2^M^ (Map C, middle) and UAP56–TREX-2^M^ (Map D, right) with the respective refined coordinate models using phenix.mtriage.

|  | THO–UAP56<br>tetramer<br>(Map A) | THO–UAP56–<br>ALYREF N-UBM<br>(Map B) | TREX2<sup>M</sup><br>(Map C) | UAP56–TREX2<sup>M</sup><br>(Map D) | UAP56 RecA2<br>–TREX2<sup>M</sup><br>(Map E) |
| --- | --- | --- | --- | --- | --- |
| Data collection |  |  |  |  |  |
| Particles | 204,147 | 204,147 | 57,499 | 37,692 | 7,741 |
| Pixel Size (Å) | 0.84 | 0.84 | 0.749 | 0.749 | 0.749 |
| Defocus range (μm) | –0.75 to –1.25 | –0.75 to –1.25 | –1.0 to –2.5 | –1.0 to –2.5 | –1.0 to –2.5 |
| Voltage (kV) | 300 | 300 | 300 | 300 | 300 |
| Electron dose (e<sup>–</sup>/Å<sup>2</sup>) | 60 | 60 | 50 | 50 | 50 |
| Reconstruction (cryoSPARC) |  |  |  |  |  |
| Resolution (Å) | 5.9 | 4.1 | 3.5 | 3.5 | 4.2 |
| Map sharpening B-factor (Å<sup>2</sup>) | –409 | –178 | –128 | –108 | –98 |
| Model composition |  |  |  |  |  |
| Non-hydrogen atoms |  | 14,961 | 6,042 | 7,984 |  |
| Protein residues |  | 1,622 | 754 | 1,012 |  |
| Nucleotide residues |  | 0 | 0 | 0 |  |
| Ligands |  | 0 | 0 | 1 |  |
| Refinement (PHENIX) |  |  |  |  |  |
| Map CC (around atoms) |  | 0.68 | 0.73 | 0.71 |  |
| B-factor (overall) |  | 146 | 125 | 126 |  |
| Rms deviations |  |  |  |  |  |
| Bond lengths (Å) |  | 0.01 | 0.01 | 0.01 |  |
| Bond angles (°) |  | 1.62 | 1.38 | 1.54 |  |
| Validation |  |  |  |  |  |
| Molprobrity score |  | 1.76 | 1.68 | 1.85 |  |
| All-atom clashscore |  | 5.63 | 4.86 | 4.72 |  |
| Rotamer outliers (%) |  | 0.06 | 0.3 | 1.57 |  |
| C-beta deviations |  | 0.1 | 0.1 | 0.2 |  |
| Ramachandran plot |  |  |  |  |  |
| Outliers (%) |  | 0.11 | 0.41 | 0.71 |  |
| Allowed (%) |  | 6.97 | 6.10 | 6.57 |  |
| Favoured (%) |  | 92.92 | 93.50 | 92.73 |  |
| Data Deposition |  |  |  |  |  |
| EMDB ID | EMD-18980 | EMD-18979 | EMD-18977 | EMD-18978 | EMD-18981 |
| PDB ID |  | 8R7L | 8R7J | 8R7K |  |

**Supplementary Table 2 | Proteomics data of native UAP56 immunoprecipitation.**

**Supplementary Table 3 | Proteomics data of native TREX–mRNP disassembly assay.**

**Supplementary Table 4 | AlphaFold2 Multimer screen results for UAP56 and its putative interactors.**

**Supplementary Table 5 | Vectors and sequences.**

## REFERENCES

Adams, P.D., Afonine, P.V., Bunkóczi, G., Chen, V.B., Davis, I.W., Echols, N., Headd, J.J., Hung, L.-W., Kapral, G.J., Grosse-Kunstleve, R.W., McCoy, A.J., Moriarty, N.W., Oeffner, R., Read, R.J., Richardson, D.C., Richardson, J.S., Terwilliger, T.C., Zwart, P.H., 2010. PHENIX: a comprehensive Python-based system for macromolecular structure solution. Acta Crystallogr. D Biol. Crystallogr. 66, 213–221. 10.1107/S0907444909052925

Afonine, P.V., Poon, B.K., Read, R.J., Sobolev, O.V., Terwilliger, T.C., Urzhumtsev, A., Adams, P.D., 2018. Real-space refinement in PHENIX for cryo-EM and crystallography. Acta Cryst D 74, 531–544. 10.1107/S2059798318006551

Aibara, S., Bai, X.-C., Stewart, M., 2016. The Sac3 TPR-like region in the Saccharomyces cerevisiae TREX-2 complex is more extensive but independent of the CID region. Journal of Structural Biology 195, 316–324. 10.1016/j.jsb.2016.07.007

Baltz, A.G., Munschauer, M., Schwanhäusser, B., Vasile, A., Murakawa, Y., Schueler, M., Youngs, N., Penfold-Brown, D., Drew, K., Milek, M., Wyler, E., Bonneau, R., Selbach, M., Dieterich, C., Landthaler, M., 2012. The mRNA-Bound Proteome and Its Global Occupancy Profile on Protein-Coding Transcripts. Molecular Cell 46, 674–690. 10.1016/j.molcel.2012.05.021

Ben-Yishay, R., Mor, A., Shraga, A., Ashkenazy-Titelman, A., Kinor, N., Schwed-Gross, A., Jacob, A., Kozer, N., Kumar, P., Garini, Y., Shav-Tal, Y., 2019. Imaging within single NPCs reveals NXF1’s role in mRNA export on the cytoplasmic side of the pore. Journal of Cell Biology 218, 2962–2981. 10.1083/jcb.201901127

Bird, R.E., Hardman, K.D., Jacobson, J.W., Johnson, S., Kaufman, B.M., Lee, S.-M., Lee, T., Pope, S.H., Riordan, G.S., Whitlow, M., 1988. Single-Chain Antigen-Binding Proteins. Science 242, 423–426. 10.1126/science.3140379

Blobel, G., 1985. Gene gating: a hypothesis. Proceedings of the National Academy of Sciences 82, 8527–8529. 10.1073/pnas.82.24.8527

Bonneau, F., Basquin, J., Steigenberger, B., Schäfer, T., Schäfer, I.B., Conti, E., 2023. Nuclear mRNPs are compact particles packaged with a network of proteins promoting RNA– RNA interactions. Genes Dev. 10.1101/gad.350630.123

Cabal, G.G., Genovesio, A., Rodriguez-Navarro, S., Zimmer, C., Gadal, O., Lesne, A., Buc, H., Feuerbach-Fournier, F., Olivo-Marin, J.-C., Hurt, E.C., Nehrbass, U., 2006. SAGA interacting factors confine sub-diffusion of transcribed genes to the nuclear envelope. Nature 441, 770–773. 10.1038/nature04752

Castello, A., Fischer, B., Eichelbaum, K., Horos, R., Beckmann, B.M., Strein, C., Davey, N.E., Humphreys, D.T., Preiss, T., Steinmetz, L.M., Krijgsveld, J., Hentze, M.W., 2012. Insights into RNA Biology from an Atlas of Mammalian mRNA-Binding Proteins. Cell 149, 1393–1406. 10.1016/j.cell.2012.04.031

Certo, M.T., Ryu, B.Y., Annis, J.E., Garibov, M., Jarjour, J., Rawlings, D.J., Scharenberg, A.M., 2011. Tracking genome engineering outcome at individual DNA breakpoints. Nat Methods 8, 671–676. 10.1038/nmeth.1648

Chang, C.-T., Hautbergue, G.M., Walsh, M.J., Viphakone, N., van Dijk, T.B., Philipsen, S., Wilson, S.A., 2013. Chtop is a component of the dynamic TREX mRNA export complex. EMBO J. 32, 473–486. 10.1038/emboj.2012.342

Cheng, H., Dufu, K., Lee, C.-S., Hsu, J.L., Dias, A., Reed, R., 2006. Human mRNA Export Machinery Recruited to the 5′ End of mRNA. Cell 127, 1389–1400. 10.1016/j.cell.2006.10.044

Chi, B., Wang, Q., Wu, G., Tan, M., Wang, L., Shi, M., Chang, X., Cheng, H., 2013. Aly and THO are required for assembly of the human TREX complex and association of TREX components with the spliced mRNA. Nucleic Acids Research 41, 1294–1306. 10.1093/nar/gks1188

Dempster, J.M., Rossen, J., Kazachkova, M., Pan, J., Kugener, G., Root, D.E., Tsherniak, A., 2019. Extracting Biological Insights from the Project Achilles Genome-Scale CRISPR Screens in Cancer Cell Lines. bioRxiv 720243. 10.1101/720243

Derrer, C.P., Mancini, R., Vallotton, P., Huet, S., Weis, K., Dultz, E., 2019. The RNA export factor Mex67 functions as a mobile nucleoporin. Journal of Cell Biology 218, 3967– 3976. 10.1083/jcb.201909028

Dias, A.P., Dufu, K., Lei, H., Reed, R., 2010. A role for TREX components in the release of spliced mRNA from nuclear speckle domains. Nat Commun 1, 97. 10.1038/ncomms1103

Dimitrova, L., Valkov, E., Aibara, S., Flemming, D., McLaughlin, S.H., Hurt, E., Stewart, M., 2015. Structural Characterization of the Chaetomium thermophilum TREX-2 Complex and its Interaction with the mRNA Nuclear Export Factor Mex67:Mtr2. Structure 23, 1246–1257. 10.1016/j.str.2015.05.002

Dou, Y., Barbosa, I., Jiang, H., Iasillo, C., Molloy, K.R., Schulze, W.M., Cusack, S., Schmid, M., Le Hir, H., LaCava, J., Jensen, T.H., 2020. NCBP3 positively impacts mRNA biogenesis. Nucleic Acids Research 48, 10413–10427. 10.1093/nar/gkaa744

Dufu, K., Livingstone, M.J., Seebacher, J., Gygi, S.P., Wilson, S.A., Reed, R., 2010. ATP is required for interactions between UAP56 and two conserved mRNA export proteins, Aly and CIP29, to assemble the TREX complex. Genes Dev 24, 2043–2053. 10.1101/gad.1898610

Ellisdon, A.M., Dimitrova, L., Hurt, E., Stewart, M., 2012. Structural basis for the assembly and nucleic acid binding of the TREX-2 transcription-export complex. Nat Struct Mol Biol 19, 328–336. 10.1038/nsmb.2235

Emsley, P., Cowtan, K., 2004. Coot: model-building tools for molecular graphics. Acta Crystallogr. D Biol. Crystallogr. 60, 2126–2132. 10.1107/S0907444904019158

Emsley, P., Lohkamp, B., Scott, W.G., Cowtan, K., 2010. Features and development of Coot. Acta Cryst D 66, 486–501. 10.1107/S0907444910007493

Evans, R., O’Neill, M., Pritzel, A., Antropova, N., Senior, A., Green, T., Žídek, A., Bates, R., Blackwell, S., Yim, J., Ronneberger, O., Bodenstein, S., Zielinski, M., Bridgland, A., Potapenko, A., Cowie, A., Tunyasuvunakool, K., Jain, R., Clancy, E., Kohli, P., Jumper, J., Hassabis, D., 2022. Protein complex prediction with AlphaFold-Multimer. bioRxiv 2021.10.04.463034. 10.1101/2021.10.04.463034

Faza, M.B., Kemmler, S., Jimeno, S., González-Aguilera, C., Aguilera, A., Hurt, E., Panse, V.G., 2009. Sem1 is a functional component of the nuclear pore complex–associated messenger RNA export machinery. Journal of Cell Biology 184, 833–846. 10.1083/jcb.200810059

Fischer, T., Rodríguez-Navarro, S., Pereira, G., Rácz, A., Schiebel, E., Hurt, E., 2004. Yeast centrin Cdc31 is linked to the nuclear mRNA export machinery. Nat Cell Biol 6, 840– 848. 10.1038/ncb1163

Fischer, T., Sträßer, K., Rácz, A., Rodriguez-Navarro, S., Oppizzi, M., Ihrig, P., Lechner, J., Hurt, E., 2002. The mRNA export machinery requires the novel Sac3p–Thp1p complex to dock at the nucleoplasmic entrance of the nuclear pores. The EMBO Journal 21, 5843–5852. 10.1093/emboj/cdf590

Gatfield, D., Le Hir, H., Schmitt, C., Braun, I.C., Köcher, T., Wilm, M., Izaurralde, E., 2001. The DExH/D box protein HEL/UAP56 is essential for mRNA nuclear export in Drosophila. Curr. Biol. 11, 1716–1721.

Gebhardt, A., Habjan, M., Benda, C., Meiler, A., Haas, D.A., Hein, M.Y., Mann, A., Mann, M., Habermann, B., Pichlmair, A., 2015. mRNA export through an additional cap-binding complex consisting of NCBP1 and NCBP3. Nat Commun 6, 8192. 10.1038/ncomms9192

Germain, H., Qu, N., Cheng, Y.T., Lee, E., Huang, Y., Dong, O.X., Gannon, P., Huang, S., Ding, P., Li, Y., Sack, F., Zhang, Y., Li, X., 2010. MOS11: A New Component in the mRNA Export Pathway. PLOS Genetics 6, e1001250. 10.1371/journal.pgen.1001250

Gibson, D.G., Young, L., Chuang, R.-Y., Venter, J.C., Hutchison, C.A., Smith, H.O., 2009. Enzymatic assembly of DNA molecules up to several hundred kilobases. Nat Meth 6, 343–345. 10.1038/nmeth.1318

Goddard, T.D., Huang, C.C., Meng, E.C., Pettersen, E.F., Couch, G.S., Morris, J.H., Ferrin, T.E., 2018. UCSF ChimeraX: Meeting modern challenges in visualization and analysis. Protein Science 27, 14–25. 10.1002/pro.3235

Golovanov, A.P., Hautbergue, G.M., Tintaru, A.M., Lian, L.-Y., Wilson, S.A., 2006. The solution structure of REF2-I reveals interdomain interactions and regions involved in binding mRNA export factors and RNA. RNA 12, 1933–1948. 10.1261/rna.212106

Gordon, J.M.B., Aibara, S., Stewart, M., 2017. Structure of the Sac3 RNA-binding M-region in the Saccharomyces cerevisiae TREX-2 complex. Nucleic Acids Research 45, 5577–5585. 10.1093/nar/gkx158

Grant, R.P., Hurt, E., Neuhaus, D., Stewart, M., 2002. Structure of the C-terminal FG-nucleoporin binding domain of Tap/NXF1. Nat Struct Mol Biol 9, 247–251. 10.1038/nsb773

Gromadzka, A.M., Steckelberg, A.-L., Singh, K.K., Hofmann, K., Gehring, N.H., 2016. A short conserved motif in ALYREF directs cap-and EJC-dependent assembly of export complexes on spliced mRNAs. Nucleic Acids Res 44, 2348–2361. 10.1093/nar/gkw009

Grünwald, D., Singer, R.H., 2010. In vivo imaging of labelled endogenous β-actin mRNA during nucleocytoplasmic transport. Nature 467, 604–607. 10.1038/nature09438

Hargous, Y., Hautbergue, G.M., Tintaru, A.M., Skrisovska, L., Golovanov, A.P., Stevenin, J., Lian, L.-Y., Wilson, S.A., Allain, F.H.-T., 2006. Molecular basis of RNA recognition and TAP binding by the SR proteins SRp20 and 9G8. The EMBO Journal 25, 5126–5137. 10.1038/sj.emboj.7601385

Hautbergue, G.M., Hung, M.-L., Walsh, M.J., Snijders, A.P.L., Chang, C.-T., Jones, R., Ponting, C.P., Dickman, M.J., Wilson, S.A., 2009. UIF, a New mRNA Export Adaptor that Works Together with REF/ALY, Requires FACT for Recruitment to mRNA. Current Biology 19, 1918–1924. 10.1016/j.cub.2009.09.041

Heath, C.G., Viphakone, N., Wilson, S.A., 2016. The role of TREX in gene expression and disease. Biochem. J. 473, 2911–2935. 10.1042/BCJ20160010

Huertas, P., Aguilera, A., 2003. Cotranscriptionally Formed DNA:RNA Hybrids Mediate Transcription Elongation Impairment and Transcription-Associated Recombination. Molecular Cell 12, 711–721. 10.1016/j.molcel.2003.08.010

Jani, D., Lutz, S., Hurt, E., Laskey, R.A., Stewart, M., Wickramasinghe, V.O., 2012. Functional and structural characterization of the mammalian TREX-2 complex that links transcription with nuclear messenger RNA export. Nucleic Acids Res 40, 4562–4573. 10.1093/nar/gks059

Jani, D., Lutz, S., Marshall, N.J., Fischer, T., Köhler, A., Ellisdon, A.M., Hurt, E., Stewart, M., 2009. Sus1, Cdc31, and the Sac3 CID Region Form a Conserved Interaction Platform that Promotes Nuclear Pore Association and mRNA Export. Molecular Cell 33, 727–737. 10.1016/j.molcel.2009.01.033

Jimeno, S., Luna, R., García-Rubio, M., Aguilera, A., 2006. Tho1, a Novel hnRNP, and Sub2 Provide Alternative Pathways for mRNP Biogenesis in Yeast THO Mutants. Mol Cell Biol 26, 4387–4398. 10.1128/MCB.00234-06

Jumper, J., Evans, R., Pritzel, A., Green, T., Figurnov, M., Ronneberger, O., Tunyasuvunakool, K., Bates, R., Žídek, A., Potapenko, A., Bridgland, A., Meyer, C., Kohl, S.A.A., Ballard, A.J., Cowie, A., Romera-Paredes, B., Nikolov, S., Jain, R., Adler, J., Back, T., Petersen, S., Reiman, D., Clancy, E., Zielinski, M., Steinegger, M., Pacholska, M., Berghammer, T., Bodenstein, S., Silver, D., Vinyals, O., Senior, A.W., Kavukcuoglu, K., Kohli, P., Hassabis, D., 2021. Highly accurate protein structure prediction with AlphaFold. Nature 596, 583–589. 10.1038/s41586-021-03819-2

Kang, G.J., Park, M.K., Byun, H.J., Kim, H.J., Kim, E.J., Yu, L., Kim, B., Shim, J.G., Lee, H., Lee, C.H., 2020. SARNP, a participant in mRNA splicing and export, negatively regulates E-cadherin expression via interaction with pinin. Journal of Cellular Physiology 235, 1543–1555. 10.1002/jcp.29073

Katahira, J., Sträßer, K., Podtelejnikov, A., Mann, M., Jung, J.U., Hurt, E., 1999. The Mex67p- mediated nuclear mRNA export pathway is conserved from yeast to human. The EMBO Journal 18, 2593–2609. 10.1093/emboj/18.9.2593

Kelly, S.M., Corbett, A.H., 2009. Messenger RNA Export from the Nucleus: A Series of Molecular Wardrobe Changes. Traffic 10, 1199–1208. 10.1111/j.1600-0854.2009.00944.x

Khong, A., Parker, R., 2020. The landscape of eukaryotic mRNPs. RNA 26, 229–239. 10.1261/rna.073601.119

Koch, B., Nijmeijer, B., Kueblbeck, M., Cai, Y., Walther, N., Ellenberg, J., 2018. Generation and validation of homozygous fluorescent knock-in cells using CRISPR–Cas9 genome editing. Nat Protoc 13, 1465–1487. 10.1038/nprot.2018.042

Köhler, A., Hurt, E., 2007. Exporting RNA from the nucleus to the cytoplasm. Nat Rev Mol Cell Biol 8, 761–773. 10.1038/nrm2255

Kozma, P., Hamori, A., Cottier, K., Kurunczi, S., Horvath, R., 2009. Grating coupled interferometry for optical sensing. Appl. Phys. B 97, 5–8. 10.1007/s00340-009-3719-1

Kurshakova, M.M., Krasnov, A.N., Kopytova, D.V., Shidlovskii, Y.V., Nikolenko, J.V., Nabirochkina, E.N., Spehner, D., Schultz, P., Tora, L., Georgieva, S.G., 2007. SAGA and a novel Drosophila export complex anchor efficient transcription and mRNA export to NPC. The EMBO Journal 26, 4956–4965. 10.1038/sj.emboj.7601901

Le Hir, H., Izaurralde, E., Maquat, L.E., Moore, M.J., 2000. The spliceosome deposits multiple proteins 20–24 nucleotides upstream of mRNA exon–exon junctions. The EMBO Journal 19, 6860–6869. 10.1093/emboj/19.24.6860

Lee, E.S., Wolf, E.J., Ihn, S.S.J., Smith, H.W., Emili, A., Palazzo, A.F., 2020. TPR is required for the efficient nuclear export of mRNAs and lncRNAs from short and intron-poor genes. Nucleic Acids Research 48, 11645–11663. 10.1093/nar/gkaa919

Lu, Q., Tang, X., Tian, G., Wang, F., Liu, K., Nguyen, V., Kohalmi, S.E., Keller, W.A., Tsang, E.W.T., Harada, J.J., Rothstein, S.J., Cui, Y., 2010. Arabidopsis homolog of the yeast TREX-2 mRNA export complex: components and anchoring nucleoporin. The Plant Journal 61, 259–270. 10.1111/j.1365-313X.2009.04048.x

Luo, M.-J., Zhou, Z., Magni, K., Christoforides, C., Rappsilber, J., Mann, M., Reed, R., 2001. Pre-mRNA splicing and mRNA export linked by direct interactions between UAP56 and Aly. Nature 413, 644–647. 10.1038/35098106

Ma, J., Liu, Z., Michelotti, N., Pitchiaya, S., Veerapaneni, R., Androsavich, J.R., Walter, N.G., Yang, W., 2013. High-resolution three-dimensional mapping of mRNA export through the nuclear pore. Nat Commun 4, 2414. 10.1038/ncomms3414

Metkar, M., Ozadam, H., Lajoie, B.R., Imakaev, M., Mirny, L.A., Dekker, J., Moore, M.J., 2018. Higher-Order Organization Principles of Pre-translational mRNPs. Molecular Cell 72, 715–726.e3. 10.1016/j.molcel.2018.09.012

Ming, Q., Gonzalez-Perez, D., Luca, V.C., 2019. Molecular engineering strategies for visualizing low-affinity protein complexes. Exp Biol Med (Maywood) 244, 1559–1567. 10.1177/1535370219855401

Mirdita, M., Schütze, K., Moriwaki, Y., Heo, L., Ovchinnikov, S., Steinegger, M., 2022. ColabFold: making protein folding accessible to all. Nat Methods 19, 679–682. 10.1038/s41592-022-01488-1

Monaco, G., Chen, H., Poidinger, M., Chen, J., de Magalhães, J.P., Larbi, A., 2016. flowAI: automatic and interactive anomaly discerning tools for flow cytometry data. Bioinformatics 32, 2473–2480. 10.1093/bioinformatics/btw191

Montpetit, B., Seeliger, M.A., Weis, K., 2012. Analysis of DEAD-Box Proteins in mRNA Export, in: Jankowsky, E. (Ed.), Methods in Enzymology, RNA Helicases. Academic Press, pp. 239–254. 10.1016/B978-0-12-396546-2.00011-5

Mor, A., Suliman, S., Ben-Yishay, R., Yunger, S., Brody, Y., Shav-Tal, Y., 2010. Dynamics of single mRNP nucleocytoplasmic transport and export through the nuclear pore in living cells. Nat Cell Biol 12, 543–552. 10.1038/ncb2056

Muhar, M., Ebert, A., Neumann, T., Umkehrer, C., Jude, J., Wieshofer, C., Rescheneder, P., Lipp, J.J., Herzog, V.A., Reichholf, B., Cisneros, D.A., Hoffmann, T., Schlapansky, M.F., Bhat, P., von Haeseler, A., Köcher, T., Obenauf, A.C., Popow, J., Ameres, S.L., Zuber, J., 2018. SLAM-seq defines direct gene-regulatory functions of the BRD4-MYC axis. Science 360, 800–805. 10.1126/science.aao2793

Ohno, M., Segref, A., Bachi, A., Wilm, M., Mattaj, I.W., 2000. PHAX, a Mediator of U snRNA Nuclear Export Whose Activity Is Regulated by Phosphorylation. Cell 101, 187–198. 10.1016/S0092-8674(00)80829-6

Pacheco-Fiallos, B., Vorländer, M.K., Riabov-Bassat, D., Fin, L., O’Reilly, F.J., Ayala, F.I., Schellhaas, U., Rappsilber, J., Plaschka, C., 2023. mRNA recognition and packaging by the human transcription–export complex. Nature 616, 828–835. 10.1038/s41586-023-05904-0

Pettersen, E.F., Goddard, T.D., Huang, C.C., Meng, E.C., Couch, G.S., Croll, T.I., Morris, J.H., Ferrin, T.E., 2021. UCSF ChimeraX: Structure visualization for researchers, educators, and developers. Protein Science 30, 70–82. 10.1002/pro.3943

Piruat, J.I., Aguilera, A., 1998. A novel yeast gene, THO2, is involved in RNA pol II transcription and provides new evidence for transcriptional elongation-associated recombination. The EMBO Journal 17, 4859–4872. 10.1093/emboj/17.16.4859

Pühringer, T., Hohmann, U., Fin, L., Pacheco-Fiallos, B., Schellhaas, U., Brennecke, J., Plaschka, C., 2020. Structure of the human core transcription-export complex reveals a hub for multivalent interactions. eLife 9, e61503. 10.7554/eLife.61503

Punjani, A., Rubinstein, J.L., Fleet, D.J., Brubaker, M.A., 2017. cryoSPARC: algorithms for rapid unsupervised cryo-EM structure determination. Nat Methods 14, 290–296. 10.1038/nmeth.4169

Ren, Y., Schmiege, P., Blobel, G., 2017. Structural and biochemical analyses of the DEAD-box ATPase Sub2 in association with THO or Yra1. eLife 6, e20070. 10.7554/eLife.20070

Richardson, C.J., Bröenstrup, M., Fingar, D.C., Jülich, K., Ballif, B.A., Gygi, S., Blenis, J., 2004. SKAR Is a Specific Target of S6 Kinase 1 in Cell Growth Control. Current Biology 14, 1540–1549. 10.1016/j.cub.2004.08.061

Rodríguez-Navarro, S., Fischer, T., Luo, M.-J., Antúnez, O., Brettschneider, S., Lechner, J., Pérez-Ortín, J.E., Reed, R., Hurt, E., 2004. Sus1, a Functional Component of the SAGA Histone Acetylase Complex and the Nuclear Pore-Associated mRNA Export Machinery. Cell 116, 75–86. 10.1016/S0092-8674(03)01025-0

Rondón, A.G., Jimeno, S., Aguilera, A., 2010. The interface between transcription and mRNP export: From THO to THSC/TREX-2. Biochimica et Biophysica Acta (BBA) - Gene Regulatory Mechanisms 1799, 533–538. 10.1016/j.bbagrm.2010.06.002

Saguez, C., Gonzales, F.A., Schmid, M., Bøggild, A., Latrick, C.M., Malagon, F., Putnam, A., Sanderson, L., Jankowsky, E., Brodersen, D.E., Jensen, T.H., 2013. Mutational analysis of the yeast RNA helicase Sub2p reveals conserved domains required for growth, mRNA export, and genomic stability. RNA 19, 1363–1371. 10.1261/rna.040048.113

Scheres, S.H.W., 2012. RELION: Implementation of a Bayesian approach to cryo-EM structure determination. Journal of Structural Biology 180, 519–530. 10.1016/j.jsb.2012.09.006

Schindelin, J., Arganda-Carreras, I., Frise, E., Kaynig, V., Longair, M., Pietzsch, T., Preibisch, S., Rueden, C., Saalfeld, S., Schmid, B., Tinevez, J.-Y., White, D.J., Hartenstein, V., Eliceiri, K., Tomancak, P., Cardona, A., 2012. Fiji: an open-source platform for biological-image analysis. Nat. Methods 9, 676–682. 10.1038/nmeth.2019

Schmidt, U., Im, K.-B., Benzing, C., Janjetovic, S., Rippe, K., Lichter, P., Wachsmuth, M., 2009. Assembly and mobility of exon-exon junction complexes in living cells. RNA 15, 862– 876. 10.1261/rna.1387009

Schneider, M., Hellerschmied, D., Schubert, T., Amlacher, S., Vinayachandran, V., Reja, R., Pugh, B.F., Clausen, T., Köhler, A., 2015. The Nuclear Pore-Associated TREX-2 Complex Employs Mediator to Regulate Gene Expression. Cell 162, 1016–1028. 10.1016/j.cell.2015.07.059

Schuller, S.K., Schuller, J.M., Prabu, J.R., Baumgärtner, M., Bonneau, F., Basquin, J., Conti, E., 2020. Structural insights into the nucleic acid remodeling mechanisms of the yeast THO-Sub2 complex. eLife 9, e61467. 10.7554/eLife.61467

Silla, T., Schmid, M., Dou, Y., Garland, W., Milek, M., Imami, K., Johnsen, D., Polak, P., Andersen, J.S., Selbach, M., Landthaler, M., Jensen, T.H., 2020. The human ZC3H3 and RBM26/27 proteins are critical for PAXT-mediated nuclear RNA decay. Nucleic Acids Research 48, 2518–2530. 10.1093/nar/gkz1238

Singh, G., Pratt, G., Yeo, G.W., Moore, M.J., 2015. The Clothes Make the mRNA: Past and Present Trends in mRNP Fashion. Annual Review of Biochemistry 84, 325–354. 10.1146/annurev-biochem-080111-092106

Skoglund, U., Andersson, K., Strandberg, B., Daneholt, B., 1986. Three-dimensional structure of a specific pre-messenger RNP particle established by electron microscope tomography. Nature 319, 560–564. 10.1038/319560a0

Smith, C., Lari, A., Derrer, C.P., Ouwehand, A., Rossouw, A., Huisman, M., Dange, T., Hopman, M., Joseph, A., Zenklusen, D., Weis, K., Grunwald, D., Montpetit, B., 2015. In vivo single-particle imaging of nuclear mRNA export in budding yeast demonstrates an essential role for Mex67p. J Cell Biol 211, 1121–1130. 10.1083/jcb.201503135

Sørensen, B.B., Ehrnsberger, H.F., Esposito, S., Pfab, A., Bruckmann, A., Hauptmann, J., Meister, G., Merkl, R., Schubert, T., Längst, G., Melzer, M., Grasser, M., Grasser, K.D., 2017. The Arabidopsis THO/TREX component TEX1 functionally interacts with MOS11 and modulates mRNA export and alternative splicing events. Plant Mol Biol 93, 283–298. 10.1007/s11103-016-0561-9

Stewart, S.A., Dykxhoorn, D.M., Palliser, D., Mizuno, H., Yu, E.Y., An, D.S., Sabatini, D.M., Chen, I.S.Y., Hahn, W.C., Sharp, P.A., Weinberg, R.A., Novina, C.D., 2003. Lentivirus-delivered stable gene silencing by RNAi in primary cells. RNA 9, 493–501. 10.1261/rna.2192803

Sträßer, K., Baßler, J., Hurt, E., 2000. Binding of the Mex67p/Mtr2p Heterodimer to Fxfg, Glfg, and Fg Repeat Nucleoporins Is Essential for Nuclear mRNA Export. Journal of Cell Biology 150, 695–706. 10.1083/jcb.150.4.695

Sträßer, K., Hurt, E., 2001. Splicing factor Sub2p is required for nuclear mRNA export through its interaction with Yra1p. Nature 413, 648–652. 10.1038/35098113

Sträßer, K., Masuda, S., Mason, P., Pfannstiel, J., Oppizzi, M., Rodriguez-Navarro, S., Rondón, A.G., Aguilera, A., Struhl, K., Reed, R., Hurt, E., 2002. TREX is a conserved complex coupling transcription with messenger RNA export. Nature 417, 304–308. 10.1038/nature746

Suzuki, K., Bose, P., Leong-Quong, R.Y., Fujita, D.J., Riabowol, K., 2010. REAP: A two minute cell fractionation method. BMC Res Notes 3, 294. 10.1186/1756-0500-3-294

Tegunov, D., Cramer, P., 2019. Real-time cryo-electron microscopy data preprocessing with Warp. Nat Methods 16, 1146–1152. 10.1038/s41592-019-0580-y

Umlauf, D., Bonnet, J., Waharte, F., Fournier, M., Stierle, M., Fischer, B., Brino, L., Devys, D., Tora, L., 2013. The human TREX-2 complex is stably associated with the nuclear pore basket. J Cell Sci 126, 2656–2667. 10.1242/jcs.118000

Viphakone, N., Cumberbatch, M.G., Livingstone, M.J., Heath, P.R., Dickman, M.J., Catto, J.W., Wilson, S.A., 2015. Luzp4 defines a new mRNA export pathway in cancer cells. Nucleic Acids Res 43, 2353–2366. 10.1093/nar/gkv070

Viphakone, N., Hautbergue, G.M., Walsh, M., Chang, C.-T., Holland, A., Folco, E.G., Reed, R., Wilson, S.A., 2012. TREX exposes the RNA binding domain of Nxf1 to enable mRNA export. Nat Commun 3, 1006. 10.1038/ncomms2005

Vorländer, M.K., Pacheco-Fiallos, B., Plaschka, C., 2022. Structural basis of mRNA maturation: Time to put it together. Curr Opin Struct Biol 75, 102431. 10.1016/j.sbi.2022.102431

Weis, K., 2002. Nucleocytoplasmic transport: cargo trafficking across the border. Current Opinion in Cell Biology 14, 328–335. 10.1016/S0955-0674(02)00337-X

Wickramasinghe, V.O., McMurtrie, P.I.A., Mills, A.D., Takei, Y., Penrhyn-Lowe, S., Amagase, Y., Main, S., Marr, J., Stewart, M., Laskey, R.A., 2010. mRNA Export from Mammalian Cell Nuclei Is Dependent on GANP. Current Biology 20, 25–31. 10.1016/j.cub.2009.10.078

Wilmes, G.M., Bergkessel, M., Bandyopadhyay, S., Shales, M., Braberg, H., Cagney, G., Collins, S.R., Whitworth, G.B., Kress, T.L., Weissman, J.S., Ideker, T., Guthrie, C., Krogan, N.J., 2008. A Genetic Interaction Map of RNA Processing Factors Reveals Links Between Sem1/Dss1-Containing Complexes and mRNA Export and Splicing. Mol Cell 32, 735– 746. 10.1016/j.molcel.2008.11.012

Xie, Y., Clarke, B.P., Kim, Y.J., Ivey, A.L., Hill, P.S., Shi, Y., Ren, Y., 2021. Cryo-EM structure of the yeast TREX complex and coordination with the SR-like protein Gbp2. eLife 10, e65699. 10.7554/eLife.65699

Xie, Y., Gao, S., Zhang, K., Bhat, P., Clarke, B.P., Batten, K., Mei, M., Gazzara, M., Shay, J.W., Lynch, K.W., Angelos, A.E., Hill, P.S., Ivey, A.L., Fontoura, B.M.A., Ren, Y., 2023. Structural basis for high-order complex of SARNP and DDX39B to facilitate mRNP assembly. Cell Reports 42, 112988. 10.1016/j.celrep.2023.112988

Zenklusen, D., Vinciguerra, P., Wyss, J.-C., Stutz, F., 2002. Stable mRNP Formation and Export Require Cotranscriptional Recruitment of the mRNA Export Factors Yra1p and Sub2p by Hpr1p. Molecular and Cellular Biology 22, 8241–8253. 10.1128/MCB.22.23.8241-8253.2002

